# Editing the core region in HPFH deletions alters fetal and adult globin expression for treatment of β-hemoglobinopathies

**DOI:** 10.1101/2022.07.11.499646

**Authors:** Vigneshwaran Venkatesan, Abisha Crystal Christopher, Prathibha Babu, Manoj Kumar K Azhagiri, Kaivalya Walavalkar, Bharath Saravanan, Saranya Srinivasan, Karthik V Karuppusamy, Annlin Jacob, Sumathi Rangaraj, Abhirup Bagchi, Aswin Anand Pai, Yukio Nakamura, Poonkuzhali Balasubramanian, Rekha Pai, Srujan Kumar Marepally, Kumarasamypet Murugesan Mohankumar, Shaji R Velayudhan, Dimple Notani, Alok Srivastava, Saravanabhavan Thangavel

## Abstract

Reactivation of fetal hemoglobin (HbF) is the commonly adapted strategy to ameliorate β-hemoglobinopathies. However, the continued production of defective adult hemoglobin (HbA) limits the HbF tetramer production affecting the therapeutic benefits. Here, we tested various deletional hereditary persistence of fetal hemoglobin (HPFH) mutations and identified a 11 kb sequence, encompassing Putative Repressor Region (PRR) to β-globin Exon-1 (βE1), as the core deletion that ablates HbA and exhibit superior production of HbF compared to HPFH or other well-established targets. The PRR-βE1 edited hematopoietic stem and progenitor cells (HSPCs) retained engraftment potential to repopulate for long-term hematopoiesis in immunocompromised mice generating HbF+ cells in vivo. Importantly, the editing induces therapeutically relevant levels of HbF to reverse the phenotypes of both sickle cell disease and β-thalassemia major. These results indicate that the PRR-βE1 gene editing in patient HSPCs can potentially lead to superior therapeutic outcomes for β-hemoglobinopathies gene therapy.

## Introduction

β-hemoglobinopathies - β-thalassemia and sickle cell disease (SCD), are highly prevalent inherited recessive globin chain disorders accounting for 3.4% of mortalities in children under the age of 5 years(*1*). β-thalassemia is caused by over 300 different mutations in the β-globin gene or its flanking nucleotides, that impair the synthesis of β-globin chain, affecting the tightly coordinated adult haemoglobin (HbA/α_2_β_2_) chain equilibrium(*2*). The excess free α-globin precipitates in erythroblasts and induces apoptosis resulting in ineffective erythropoiesis(*3*). SCD is caused by the E6V (rs334) missense mutation in the β-globin gene that causes polymerization of deoxygenated sickle hemoglobin (HbS) tetramers, which severely reduces their circulating lifespan, and eventually causes vascular damage and progressive multiorgan damage(*4*).

Morbidity in β-thalassemia and SCD patients is inversely correlated with the levels of fetal haemoglobin (HbF) in adulthood(*5, 6*). Expression of γ-globin, the fetal β-like globin component of HbF, improves the globin chain equilibrium and thus prevents apoptosis of erythroid cells in β-thalassemia. Similarly, in SCD, γ-globin competes with the sickle β-globin chains (βs) to form HbF tetramer (α_2_γ_2_), thereby reducing the production of sickle RBCs. Hence, much attention has focused on identifying and manipulating genetic factors involved in HbF regulation(*7–9*). Two recent clinical studies involving shRNA mediated erythroid specific downregulation of BCL11A and gene editing-mediated disruption of its erythroid-specific enhancer have demonstrated reactivated HbF levels sufficient to reach transfusion independence(*10, 11*). However, up to 50% of hemoglobin remained as HbS in the SCD patients, suggesting the need to inhibit defective β-globin production. Hereditary persistence of fetal haemoglobin (HPFH) mutations are documented to produce varying levels of HbF in healthy individuals without any deleterious effects(*12*). Importantly, the HPFH mutations are beneficial in alleviating disease severity when co-inherited with β-hemoglobinopathies(*13*). Among the genetic variants that induce HbF expression, deletional HPFH mutations produce higher levels of HbF and are highly prevalent(*14*). β-globin production is blocked in the deletional HPFH mutations, distinguishing them from other HbF reactivating mutations. HPFH deletions range in size from 12.9kb to 84.9kb, encompassing *HBG1*, *HBBP1*, *HBD* and *HBB* genes in the β-globin cluster, and result in pancellular HbF production(*15*). Studies have shown that the introduction of HPFH deletions in adult hematopoietic stem and progenitor cells (HSPCs) results in activation of γ-globin with subsequent amelioration of the sickle phenotype(*16, 17*). However, the minimal genomic deletion required for therapeutically relevant γ-globin activation, engraftment, and repopulation potential of HSPCs harboring such genomic deletions and their efficacy in reversing the disease phenotype, are yet to be revealed.

To explore the translational potential of HSPCs harbouring deletional HPFH mutations, we screened various HPFH deletions created by CRISPR-Cas9 and identified an 11kb regulatory sequence PRR-βE1. We show that HSPCs carrying the PRR-βE1 deletion retain long-term engraftment potential, activate γ-globin expression in erythroblasts to higher levels than the known HbF inducing targets, silence defective β-globin, and alleviate β-hemoglobinopathies.

## Results

### Genomic deletion encompassing PRR to exon1 of β-globin is sufficient to reproduce deletional HPFH phenotype

To identify a HPFH deletion suitable for therapeutic gene editing, we introduced deletional HPFH mutations of <30 kb in size - Algerian(*18*) (24 kb), French(*18*) (20 kb), South-East (SE) Asian(*19*) (27 kb) and Sicilian(*20*) (12.9 kb) by CRISPR-Cas9 dual gRNA gene editing in HUDEP-2 cell lines (Fig. 1A). The 7.2 kb Corfu deletion, which is now considered as δ-β thalassemia and requires homozygous deletion for activating therapeutic HbF levels(*21, 22*), was excluded from our screening.

**Figure 1.**
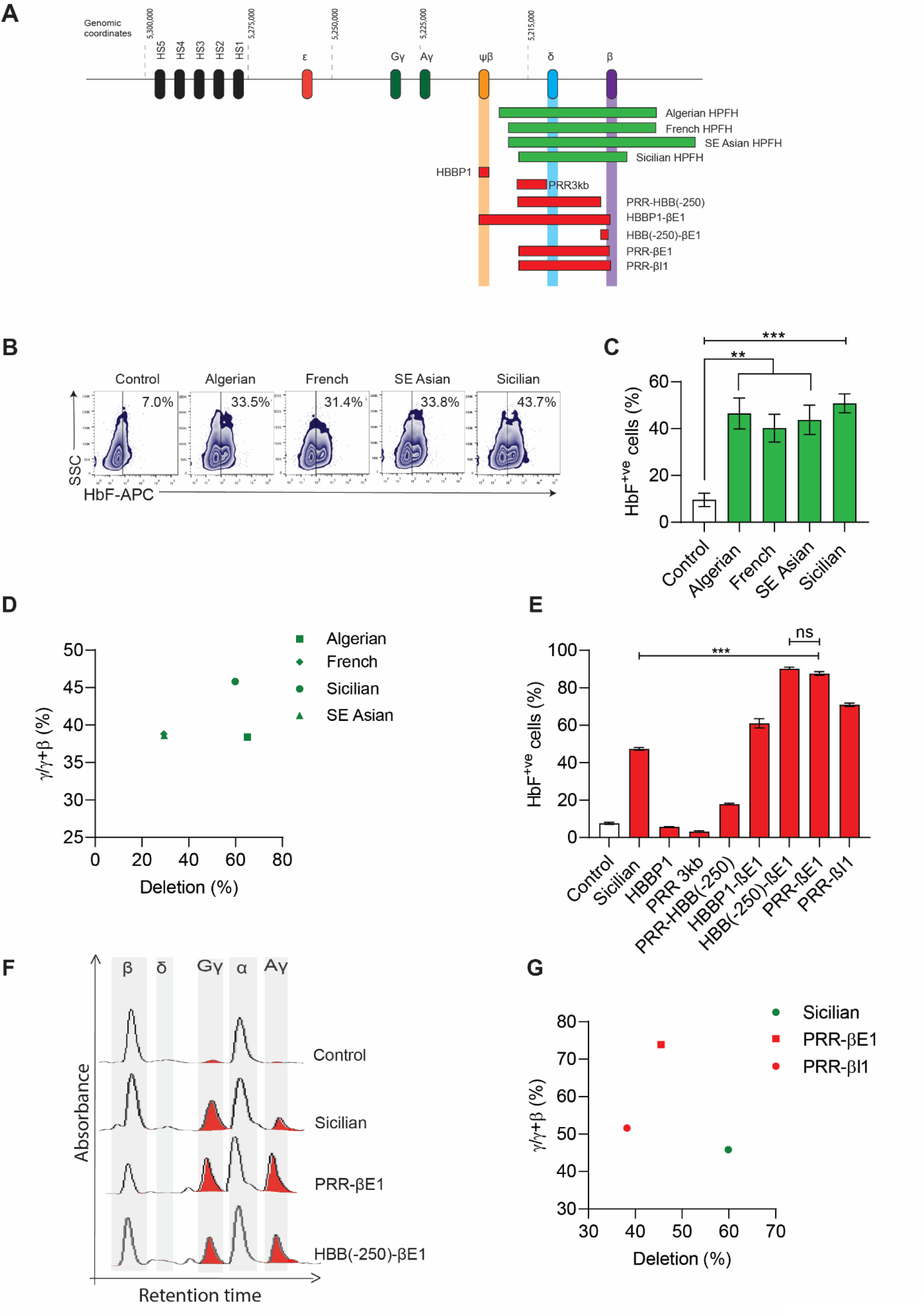
Genomic deletion encompassing PRR to exon1 of β-globin is sufficient to reproduce deletional HPFH phenotype. A. Diagrammatic representation of β-globin cluster and the break points of naturally occurring HPFH deletions (green). These deletions are introduced in the experiments shown in figure 1B-D. Break points of deletions (red) introduced in the experiment shown in figure 1E-G. All these deletions were generated in HUDEP-2 cell lines by CRISPR/Cas9 dual gRNA approach. B. Representative flow cytometry plot of HbF^+ve^ cells. HUDEP-2 cell lines gene edited for naturally occurring HPFH deletions, differentiated into erythroblasts and analysed for HbF^+ve^ cells on day 8 of erythroid differentiation. Inset shows percentage of HbF+ cells. C. Percentage of HbF^+ve^ cells upon introducing various HPFH deletions. n = 4 D. Percentage of γ/γ+β ratio and the genomic deletion. Globin chain levels was measured by HPLC chain analysis, and the deletion was quantified by ddPCR approach. The graph contains representative data of 1 of the 3 independent experiments. E. Percentage of HbF^+ve^ cells upon introducing deletions in the region encompassing Sicilian HPFH. n = 2. F. Representative globin chain HPLC chromatograms. G. Percentage of γ/γ+β ratio and the genomic deletion upon introducing deletions in the region encompassing Sicilian HPFH. Globin chain activation was measured by HPLC chain analysis, and the deletion was quantified by ddPCR approach. The graph contains representative data of 1 of the 2 independent experiments. n = 1. Error bars represent mean ± SEM, ns; non-significant. ^∗∗^p ≤ 0.01, ^∗∗∗^p ≤ 0.001 (Multiple t test).

Erythroid differentiation of gene edited HUDEP-2 cells showed an increased percentage of HbF^+ve^ cells in all of the deletions (Fig. 1B, C). Sicilian HPFH induced the highest activation of γ-globin chains (Fig. 1D, and S1A). Next, to decipher the HbF regulatory region in Sicilian HPFH deletion, we excised different regions spanning the deletion such as pseudo-β region (HBBP1), Putative Repressor Region (PRR 3kb), PRR to upstream region of β-globin promoter (PRR-HBB(-250)), promoter to exon1 (HBB(-250)-βE1), HBBP1 to exon1 (HBBP1-βE1), PRR to exon1 (PRR-βE1) and PRR to intron-1 (PRR-βI1) of the β-globin gene (Fig. 1A). Among these candidates, PRR-βE1 and HBB (-250)-βE1 showed 1.8-fold higher percentage of HbF^+ve^ cells than the Sicilian HPFH (Fig. 1E). Both these candidates had similar increase in the levels of γ-globin mRNA (Fig. S1B) and γ/γ+β ratio (Fig. S1C). However, HBB(-250)-βE1 deletion resembling the British black and Croatian β-thalassemia genotype(*23*), showed mild reduction in the proportion of mature erythroid subsets marked as CD235a^+^CD71^-^ (Fig. S1D). Therefore PRR-βE1 was considered for further studies.

The increased activation of γ-globin in PRR-βE1 edited cells over the Sicilian HPFH was reproducible on altering the βE1 cut site with a different sgRNA (Fig 1F, G). While Sicilian HPFH favored Gγ activation, the PRR-βE1 edited cells displayed equivalent activation of both Aγ and Gγ chains (Fig. S1E). Variant HPLC analysis confirmed that the activated γ paired with α chains to form a functional HbF tetramer (Fig. S1F, G). The HbF tetramer of 10% and above has demonstrated therapeutic benefits(*24*). The PRR-βE1 editing produced 79.3±3% of HbF tetramer (vs 0.2±0.1% in control) and reduced HbA tetramer to 17.8±3% (vs 98.1±0.2% in control).

We further characterised the PRR-βE1 region by gene editing only the cut sites (PRR or βE1) with single sgRNA or the entire region (PRR-βE1) by dual sgRNA to analyse the contribution of the individual cut sites in HbF activation. While PRR cut site editing alone had no effects, βE1cut site editing increased HbF^+ve^ cells and γ-globin levels and reduced β-globin levels. PRR-βE1 edited cells had a heterogenous genotype with PRR indels (11.8±1%), βE1 indels (12.6±1%) and PRR-βE1 deletion (58.4±5%) (Fig. S1H) and produced the higher percentage of HbF^+ve^ cells than editing βE1 alone (Fig. S1I). The proportion of γ-globin chains and the reduction in β-globin chains were similar to βE1 editing (Fig. S1J). The ddPCR mediated quantification of PRR-βE1 gene editing events (Fig. S2A, B) was confirmed by gap PCR analysis in the FACS sorted single cell clones. The analysis further showed that these cells had biallelic (45.4%) or monoallelic (21.2%) deletion of entire PRR-βE1 (Fig. S2C, D). The PRR-βE1 is a 11 kb region and was intact only in 9% of the wild type clones. Around 24.2% of clones had a shorter deletion (<11kb) and 3.2% of clones had deletion >11kb. The PRR-βE1 region is excised in all the deletional HPFH mutations (Fig. S3). The sequence spanning PRR is intact in δβ thalassemia and β-thalassemia deletions, suggesting it as a region that distinguishes HPFH and thalassemia phenotypes.

### Robust γ-globin induction and β-globin silencing in the erythroblasts differentiated from PRR-βE1 edited HSPCs

To probe the effect of the PRR-βE1 deletion in therapeutically relevant cells, G-CSF mobilised HSPCs from 4 different healthy donors were electroporated with Cas9 RNP targeting PRR and βE1 sites individually and in combination. The total gene editing efficiency in PRR-βE1 was 92±7%, among which PRR-βE1 deletion comprised 74.3±2% (Fig. 2A). The viability of PRR-βE1 edited HSPCs remained similar to that of AAVS1 edited cells 72hrs post-editing (Fig. S4A). On differentiation of HSPCs into erythroblasts using a three phase *in vitro* erythroid differentiation protocol, a significant increase in the percentage of HbF^+ve^ cells were observed in PRR-βE1 (78.2±3) and βE1 (58.4±7) edited cells compared to the control (26.3±1) (Fig. 2B). A 5-fold increase in the γ-globin mRNA (Fig. S4B) and up to 8.1-fold higher levels of HbF tetramers as indicated by variant HPLC analysis were observed in βE1 and PRR-βE1 editing (Fig. 2C).

**Figure 2.**
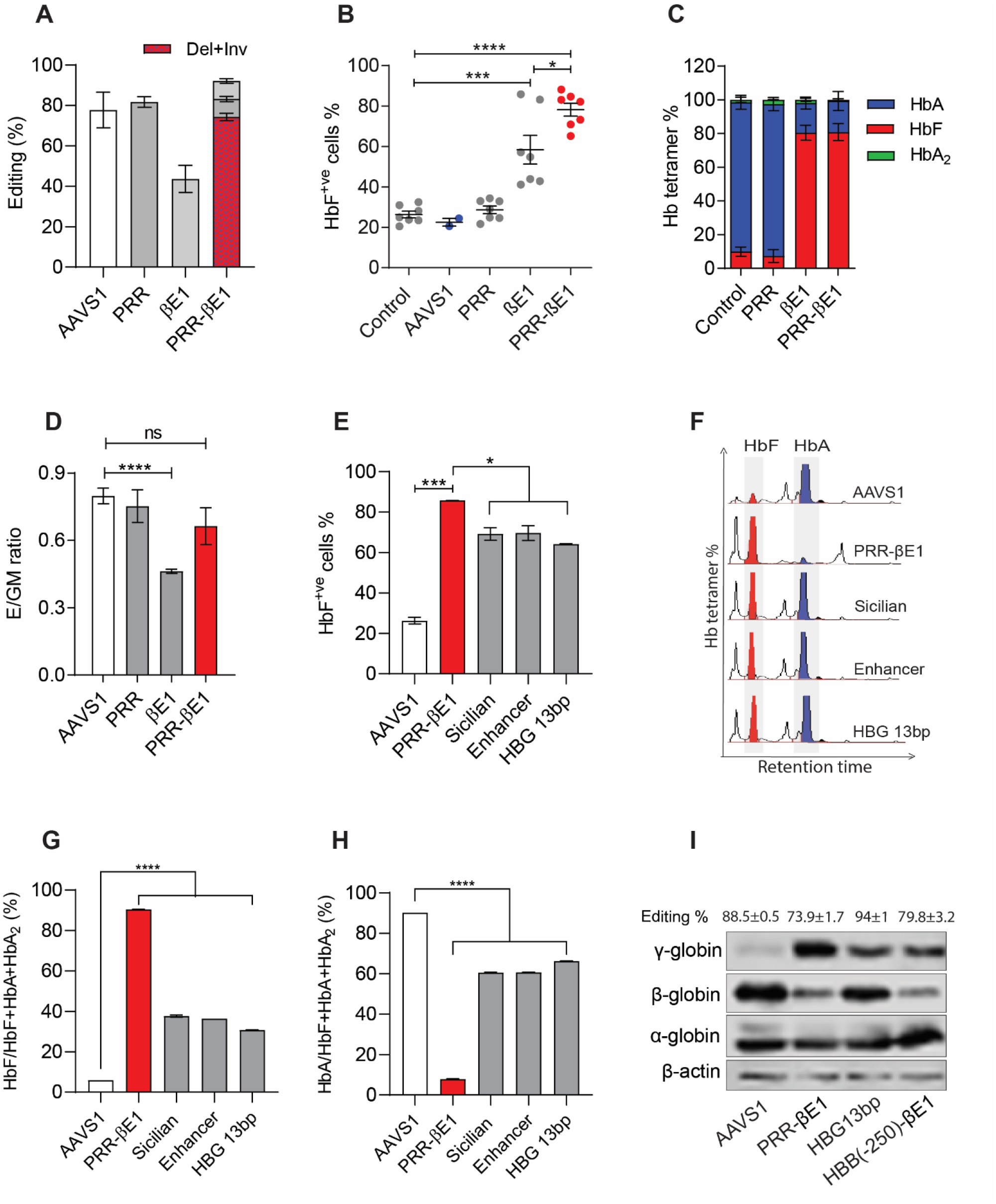
Robust γ-globin induction and β-globin silencing in the erythroblasts differentiated from PRR-βE1 edited HSPCs. A. Percentage of gene editing in PRR, βE1, and PRR-βE1 gene edited healthy donor HSPCs. Indels measured by sanger sequencing and ICE analysis. Deletion + Inversion (Del+Inv) (red checker box) in PRR-βE1 quantified by ddPCR. The PRR-βE1 edited cells had deletion, indels at PRR region and βE1. Donor = 4, n = 8 B. FACS analysis of percentage of HbF^+ve^ cells in erythroblasts generated from gene edited HSPCs. Gene editing targets are indicated at the bottom. Control refers unedited cells. Each dot indicates individual experiment. Donor = 4, n = 6. C. Proportion of hemoglobin tetramers. HSPCs were gene edited for PRR, βE1 and PRR-βE1 and differentiated into erythroblasts for variant HPLC analysis. Donor = 1, n = 2. D. Ratio of Erythroid (E) to granulocyte-monocyte (GM) CFU colonies. HSPCs were gene edited for AAVS1, PRR, βE1 and PRR-βE1 and plated in methocult medium. Both BFU-E and CFU-E colonies were considered as erythroid (E) colony. Donor = 2, n = 4. E. FACS analysis of percentage of HbF^+ve^ cells in erythroblasts generated from gene edited HSPCs. Gene editing targets are indicated at the bottom. Enhancer refers to BCL11A enhancer, HBG 13bp refers to HBG promoter. Donor = 1, n = 2 F. Representative hemoglobin variant HPLC chromatograms showing HbF and HbA tetramers in gene edited cells. G. Proportion of HbF among HbF, HbA, and HbA_2_ tetramers. HSPCs were gene edited for PRR-βE1, Sicilian HPFH, BCL11A enhancer, and HBG promoter 13bp, differentiated into erythroblasts and analysed by variant HPLC. Donor = 1, n = 2 H. Proportion of HbA among HbF, HbA, and HbA_2_ tetramers. HSPCs were gene edited for PRR-βE1, Sicilian HPFH, BCL11A enhancer, and HBG promoter 13bp, differentiated into erythroblasts and analysed by variant HPLC. Donor = 1, n = 2. I. Representative western blot image showing the band intensity of globin chains for erythroblasts derived from AAVS1, PRR-βE1, HBG13bp, and HBB(-250)-βE1 gene edited HSPCs. Donor = 1, n = 1. Error bars represent mean ± SEM, ns; non-significant. ^∗^p ≤ 0.05, ^∗∗∗^p ≤ 0.001, ^∗∗∗∗^p ≤ 0.0001 (Multiple t test).

No significant difference was observed in the frequency of editing in HSPCs and in erythroblasts derived from gene edited HSPCs (Fig. S4C). The erythroblast proliferation (Fig. S4D), and erythroid maturation analysis using CD235a and Hoechst(*25*), indicated reticulocyte generation in the PRR-βE1 edited cells at a frequency similar to the control (Fig. S4E). A modest decrease in the reticulocyte frequency on βE1 editing was noticed. Evaluation of erythroid colony forming unit (CFU) potential measured by the ratio of erythroid (BFU-E+CFU-E) to granulocyte-monocyte (GM) generation, showed a significant reduction on βE1 editing, whereas the ratio remained comparable to AAVS1 control on PRR-βE1 editing (Fig. 2D). Analyses of retention of gene editing in the erythroblasts, proliferation kinetics, CFU potential and the erythroid marker, indicate a normal erythropoiesis in PRR-βE1 edited HSPCs.

We compared the efficacy of HbF generation by PRR-βE1 and Sicilian HPFH with the well characterized targets -BCL11A erythroid specific enhancer and HBG promoter 13bp that have advanced into clinical studies(*11, 26*). Gene editing efficiency of HSPCs (Fig. S4F) and the ratio of Erythroid to GM colonies for all these targets were comparable (Fig. S4G). On erythroid differentiation, all these targets produced HbF^+^ cells and among them, the PRR-βE1 cells produced a highest percentage of HbF^+ve^ cells (Fig. 2E). The HbF tetramer production was 2-fold higher and HbA tetramer production was 7-fold lower in PRR-βE1 edited cells than the other targets (Fig. 2F-H). Western blot analysis further confirmed the increased γ-globin chains with simultaneous decrease in β-globin chains on PRR-βE1 editing (Fig. 2I, S4H).

### The PRR-βE1 deletion is retained in long-term engrafting HSCs in NSG mice

To characterise the *in vivo* reconstitution potential of PRR-βE1 gene edited cells, HSPCs from two different healthy donors were gene edited with PRR-βE1 and crRNA less RNP (control). The crRNA less RNP will not introduce any DNA double stand breaks and thus could serve as an optimal control to monitor the engraftment defects associated with genomic deletion. The edited cells were transplanted into 7-8 weeks old sub-lethally irradiated NSG mice.

On analysing human cells in mice bone marrow (BM) at 16 weeks post transplantation, percentage of engraftment was comparable in control and PRR-βE1 edited HSPCs for both the donors (Fig. 3A, Fig. S5A). The multilineage repopulation potential remained similar, indicating no lineage bias on PRR-βE1 gene editing (Fig. 3B). In vitro erythroid differentiation of engrafted cells from mouse BM showed increased HbF^+ve^ cells (Fig. 3C) and γ/(γ+β) ratio (Fig. 3D) upon PRR-βE1 gene editing. Analysis of percentage of reticulocytes (Fig. S5B) and CD235a^+^/CD71^-^ erythroblasts (Fig. S5C) confirmed an intact erythroid maturation in these cells. Genotyping analysis indicated the retention of PRR-βE1 genomic deletion in long-term repopulating cells, confirming that γ-globin activation is stimulated by PRR-βE1 deletion (Fig. 3E).

**Figure 3.**
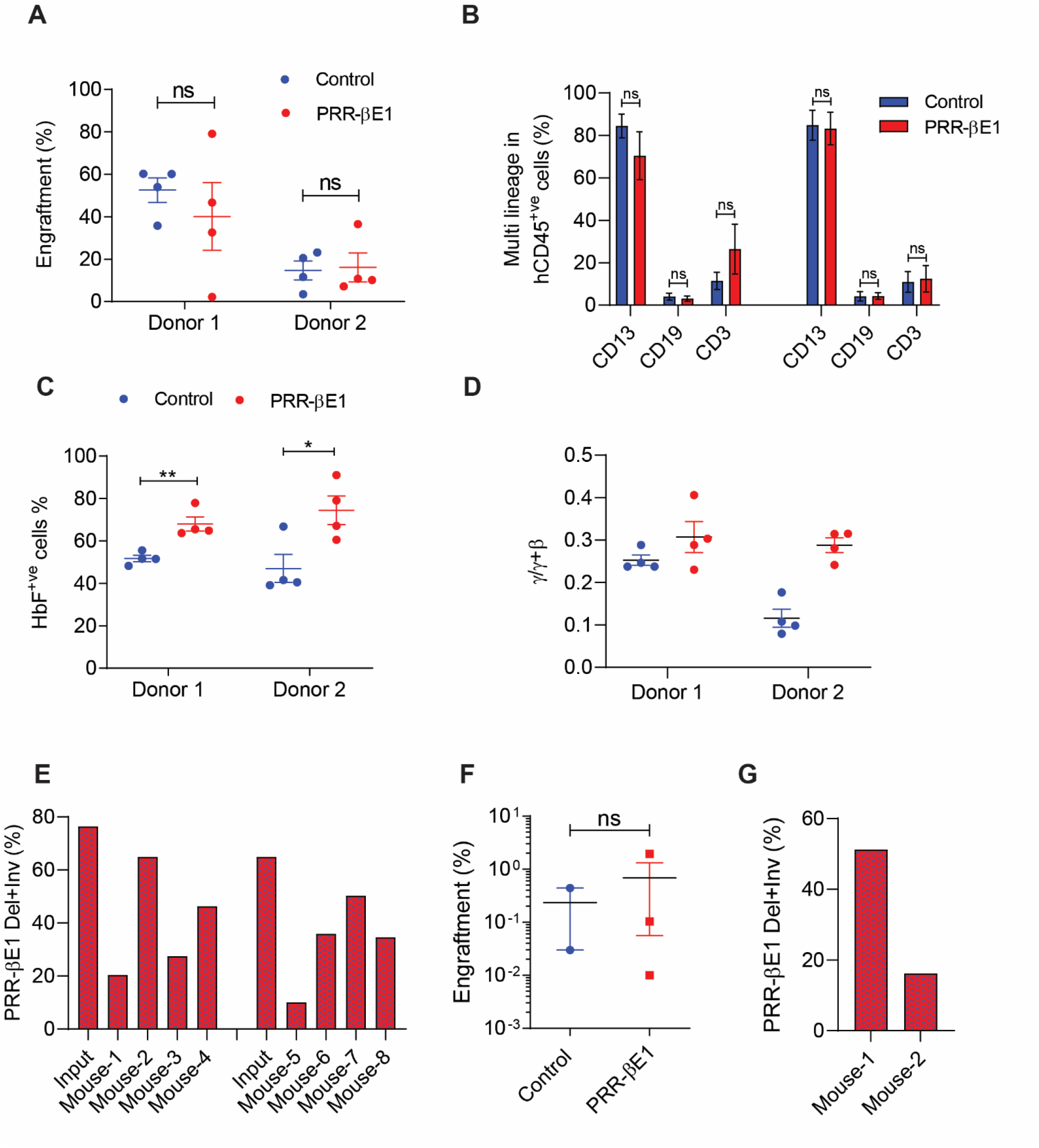
The PRR-βE1 deletion is retained in long-term engrafting HSCs in NSG mice. A. Percentage of bone marrow engraftment of control and PRR-βE1 gene edited HSPCs. Transplantation was performed in NSG mice and bone marrow was analyzed 16 weeks post transplantation. Each dot depicts individual mouse. A targeting crRNA less Cas9 RNP was used as a control. Donor = 2, n = 8. B. Percentage of lineage cells in bone marrow CD13 (monocyte), CD19 (B-cells), and CD3 (T-cells) in engrafted hCD45^+ve^ cells. C. Percentage of HbF^+ve^ cells generated on in vitro erythroid differentiation of bone marrow engrafted cells. D. HPLC chain analysis of ratio of γ/γ+β chains in the erythroblasts. The 16 weeks engrafted cells in bone marrow of NSG mice were differentiated into erythroblasts and analyzed by chain HPLC analysis. E. Percentage of PRR-βE1 deletion+inversion (Del+Inv) in PRR-βE1 gene edited HSPCs in infused fraction (input) and in engrafted cells of bone marrow, 16 weeks post transplantation. F. Secondary engraftment analysis: The long-term (16 weeks) engrafted cells in the bone marrow of NSG mice were collected and 2×10^6^ cells were infused into a secondary NSG recipients and analyzed 8 weeks post infusion. Donor = 1, n = 3. Percentage of PRR-βE1 deletion/inversion (Del+Inv) in engrafted bone marrow cells of secondary recipients as measured by ddPCR. Donor = 1, n = 2. Error bars represent mean ± SEM, ns; non-significant. ^∗^p ≤ 0.05, ^∗∗^p ≤ 0.01, ^∗∗∗^p ≤ 0.001 (unpaired t test, two-tailed).

To assess the serial repopulation potential of HSCs harbouring PRR-βE1 deletion, we infused BM cells of primary recipient to sub lethally irradiated secondary recipient. Engraftment analysis at 8 weeks post infusion showed similar frequency of engraftment in PRR-βE1 cells and in control (Fig. 3F). Analysis of PRR-βE1 deletion in the engrafted cells of secondary recipients showed retention of PRR-βE1 genomic deletion (Fig. 3G).

### The PRR-βE1 deletion in HSPCs results in increased HbF^+ve^ red cells *in vivo*

To extensively characterise engraftment characteristics of PRR-βE1 gene edited HSPCs, we employed NBSGW mouse, which supports robust human cell engraftment and erythropoiesis(*27*). HSPCs gene edited with PRR-βE1 or crRNA less RNP (control) were infused into 7-8 weeks old busulfan conditioned NBSGW mice. On 16 weeks post transplantation, the engraftment of PRR-βE1 edited cells in mice bone BM, peripheral blood, and spleen were comparable to control (Fig. 4A-C). Similarly, the multi-lineage repopulation potential of engrafted cells in the BM was also comparable between samples. The percentage of CD235a^+ve^ erythroblasts were similar, confirming the intact erythropoiesis *in vivo* (Fig. 4D).

**Figure 4.**
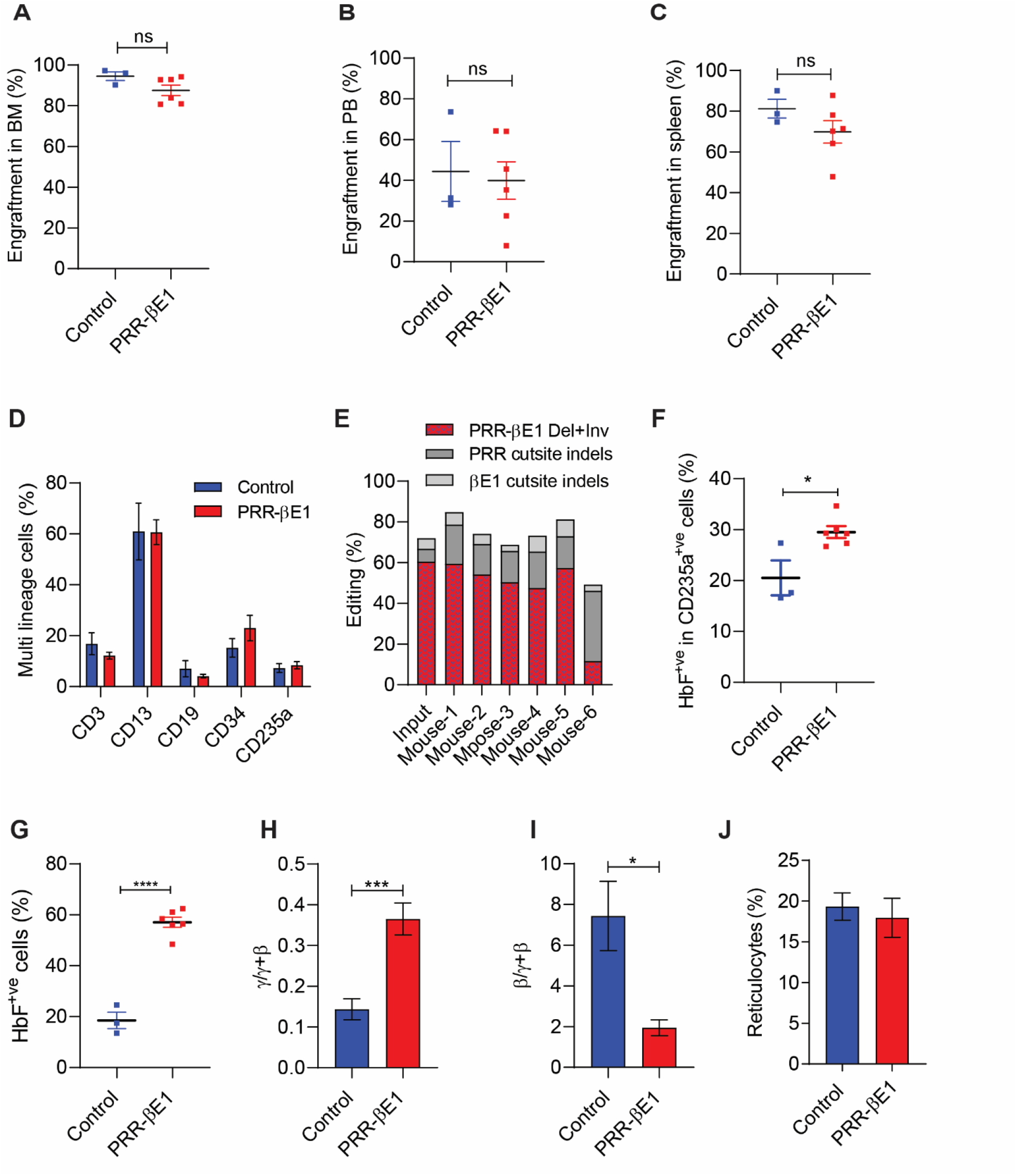
The PRR-βE1 deletion in HSPCs results in increased HbF^+ve^ red cells *in vivo*. Control and PRR-βE1 gene edited healthy donor HSPCs were transplanted into NBSGW mice and analysed 16 weeks post transplantation. Each dot indicates a single mouse. Donor = 1. Error bars represent mean ± SEM, ns; non-significant. ^∗^p ≤ 0.05, ^∗∗∗^p ≤ 0.001, ^∗∗∗∗^p ≤ 0.0001 (unpaired t test, two-tailed). A. Percentage of engraftment in the bone marrow (BM) B. Percentage of peripheral blood (PB) chimerism. C. Percentage of engraftment in the spleen. D. Percentage of lineage markers in BM – CD3 (T-cells), CD13 (monocyte), CD19 (B-cells), CD34 (HSPCs), and CD235a (Erythroid) in engrafted cells. CD235a^+^ cells were analyzed form CD45^-^ cells. E. Percentage of PRR-βE1 deletion+inversion (Del+Inv), PRR cut site indels, and βE1 cut site indels in PRR-βE1 gene edited HSPCs in infused fraction and in engrafted cells. F. Percentage of HbF^+ve^ cells in hCD235a^+ve^ cells obtained from mouse BM. G. Percentage of HbF^+ve^ cells generated by erythroid differentiation of engrafted cells in the BM. H. Ratio of γ/γ+β chains. Mouse BM was collected, *in vitro* differentiated into erythroblasts and analyzed by chain HPLC. I. Ratio of β/γ+β chains. Mouse BM was collected, *in vitro* differentiated into erythroblasts and analyzed by chain HPLC. J. Percentage of reticulocytes. Mouse BM was collected, *in vitro* differentiated into erythroblasts and analyzed by FACS.

Importantly, genotyping of the long-term repopulating cells in all the mice indicated the retention of PRR-βE1 deletion and the percentage of deletion was comparable to input cells (Fig. 4E). Next, we sorted human CD235a^+ve^ erythroblasts from mouse BM and observed a significant increase in the percentage of HbF^+ve^ cells *in vivo* on PRR-βE1 editing (Fig. 4F). Further, *in vitro* erythroid differentiation of cells retrieved from mouse BM cells showed significant increase in HbF^+ve^ cells (Fig. 4G), γ/(γ+β) ratio (Fig. 4H) and decreased β/(γ+β) ratio (Fig. 4I) with comparable reticulocyte production (Fig. 4J).

### PRR-βE1 gene edited patient HSPCs reverses sickle cell disease phenotype

To test the potential of PRR-βE1 gene editing strategy in the reversal of sickle phenotype, the plerixafor mobilized HSPCs from two SCD patients of compound heterozygous genotype HbS/CD41/CD42(-TCTT) and HbS/IVS1-5 (Fig. S6A) were gene edited with Cas9-RNP targeting AAVS1, PRR, βE1 and PRR-βE1. The gene editing frequency in each condition was > 80%, with PRR-βE1 deletion frequency of > 56% (Fig. 5A). The gene edited cells were *in vitro* differentiated into erythroblasts under hypoxia (5% O2)(*28*) and the erythroblasts derived from PRR-βE1 and βE1 gene edited HSPCs showed a significant increase in the γ-globin mRNA expression (Fig. 5B) and the percentage of HbF^+ve^ cells (Fig. 5C). Next, we performed sickling assay by treating erythroblasts with sodium metabisulfite. On treatment, AAVS1 edited cells underwent sickling while both βE1 and PRR-βE1 edited group had 12-fold (HbS/CD41/CD42(-TCTT)) and 30-fold (HbS/IVS1-5) reduction in the sickling (Fig. 5D-E). Variant HPLC analysis further showed that entire hemoglobin in the βE1 and PRR-βE1 are composed of HbF tetramers and with near complete reduction of HbS (Fig. 5F-G). These observations demonstrate the therapeutic potential of PRR-βE1 gene editing for SCD gene therapy (Fig. S9).

**Figure 5.**
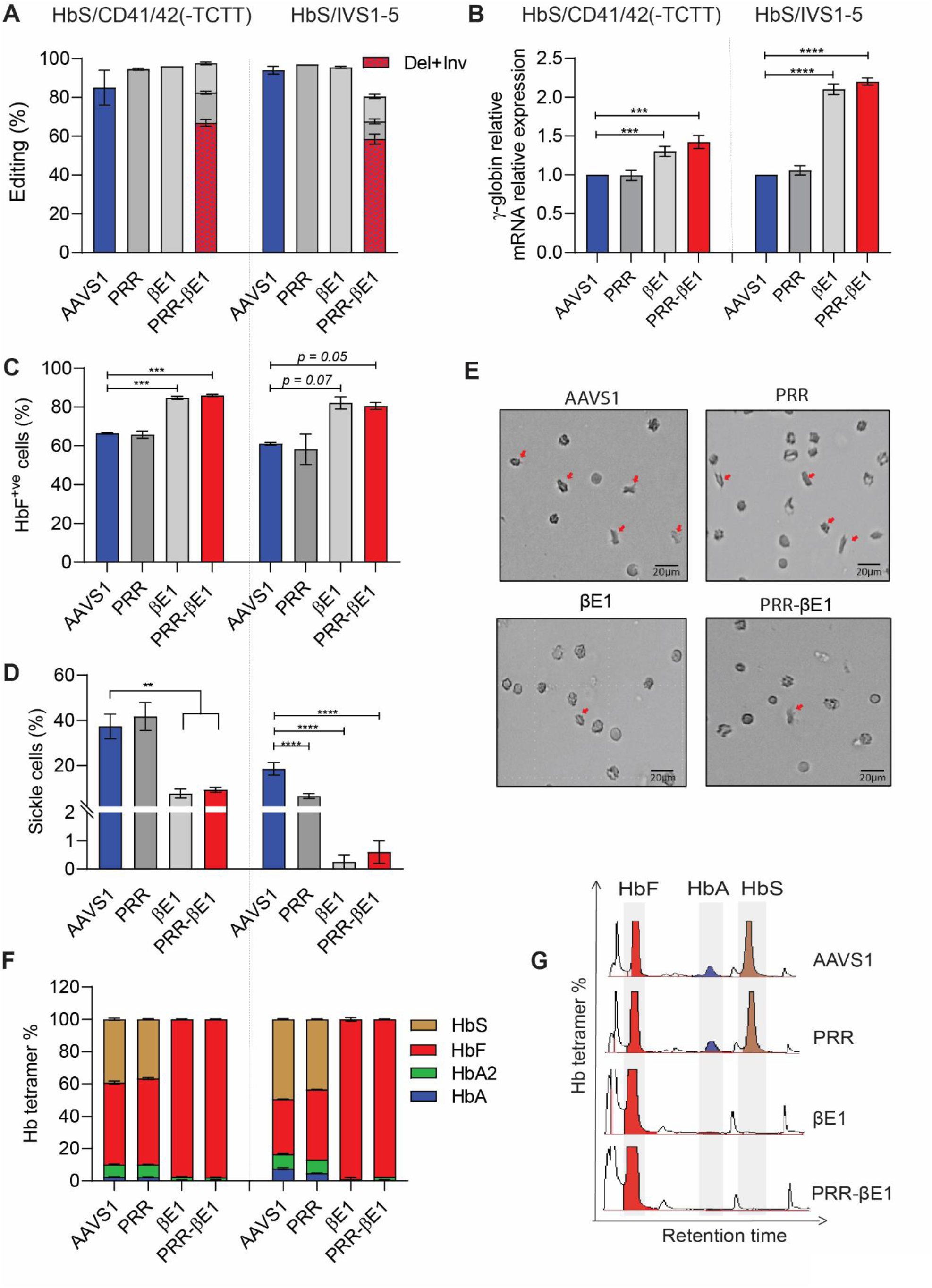
PRR-βE1 gene edited patient HSPCs reverses sickle cell disease phenotype. Plerixafor mobilized HSPCs from sickle cell patients of genotype HbS/CD41/42(-TCTT) and HbS/IVS1-5 were gene edited for AAVS1, PRR, βE1, and PRR-βE1. Error bars represent mean ± SEM. ^∗∗^p ≤ 0.01, ^∗∗∗^p ≤ 0.001, ^∗∗∗∗^p ≤ 0.0001 (Multiple t test). A. Percentage of gene editing. Indels measured by sanger sequencing and ICE analysis. Deletion/inversion (Del+Inv) (red checker box) in PRR-βE1 quantified by ddPCR. Donor = 2, n = 4. B. Relative globin mRNA expression. The patient HSPCs were gene edited for PRR, βE1, and PRR-βE1 and differentiated into erythroblasts. Real-time PCR analysis was used for mRNA quantification and the globin chain expression was normalised with β-actin. The patient genotype is indicated at the bottom. Donor = 2, n = 4. C. Percentage of HbF^+ve^ cells. The gene edited patient HSPCs were differentiated into erythroblasts and intracellular HbF positive cells were analyzed by FACS. Donor = 2, n = 4. D. Percentage of sickle cells. Gene edited patient HSPCs were differentiated into erythroblasts in hypoxia (5% O2) and treated with 1.5% sodium metabisulfite. Cells were scored from random fields using EVOS FL Auto Imaging System microscope. Atleast 8 fields were analysed. Each field contained a minimum of 150 cells. Donor = 2, n = 4. E. Representative image of sickle cells (red arrow) and non-sickled cells. F. Proportion of hemoglobin tetramer. The gene edited patient HSPCs were differentiated into erythroblasts and the hemoglobin tetramers were analyzed by variant HPLC. Donor = 2, n = 4. G. Representative variant HPLC chromatogram showing HbA, HbF and HbS. Donor = 2, n = 4.

### PRR-βE1 gene edited patient HSPCs reverse β-thalassemia phenotype

To test the therapeutic potential of our gene editing strategy in reversing β-thalassemia defects, we edited HSPCs obtained from β-thalassemia patients of three different β^0^β^0^ genotypes: CD26 (G>A)/IVS1-5 (G>C), IVS1-5 (G>C), and CD30 (G>A) (Fig. S6B). These β thalassemia mutations are highly prevalent in India and Southeast Asian countries (*30, 31*). Due to poor PBMNCs yield after mobilization, CD26 (G>A)/IVS1-5 (G>C) PBMNCs were differentiated into erythroblasts and edited on day 8 of erythroid differentiation. The PRR-βE1 gene editing efficiency remained >80% in all the genotypes (Fig. 6A). On *in vitro* erythropoiesis, βE1 and PRR-βE1 cells showed a significant increase in the frequency of HbF^+ve^ cells (Fig. S7A) and γ/(γ+β) ratio (Fig. S7B). The levels of β-globin were reduced (Fig. S7C). The ratio of α to non-α-globin chains were also observed to be reduced (Fig. 6B, C) suggesting the reduction of free α-globin chains.

**Figure 6.**
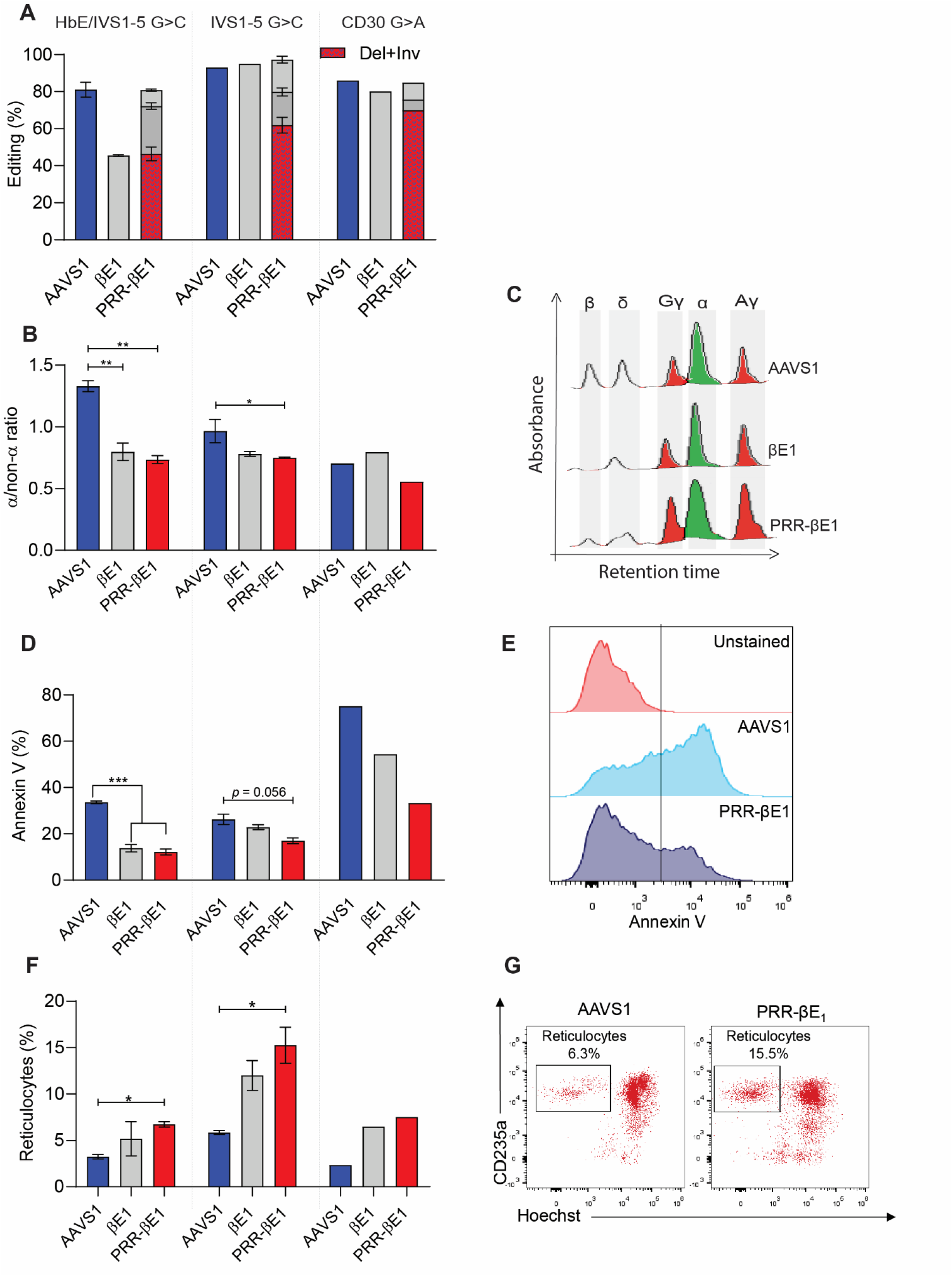
PRR-βE1 gene edited patient HSPCs reverse β-thalassemia phenotype. The G-CSF mobilized HSPCs from β-thalassemia patients of genotype IVS1-5 (G>C), and CD30 (G>A) were gene edited for AAVS1, βE1, and PRR-βE1. For HbE (G>A)/IVS1-5 (G>C), the PBMNCs were differentiated into erythroblasts and gene edited for AAVS1, βE1, and PRR-βE1. Error bars represent mean ± SEM. ^∗^p ≤ 0.05, ^∗∗^p ≤ 0.01, ^∗∗∗^p ≤ 0.001 (Multiple t test). Donor = 3, n = 6. A. Percentage of gene editing in HSPCs. Deletion/Inversion (Del+Inv) in PRR-βE1 as quantified by ddPCR. indels of the cut sites PRR and βE1as measured by ICE analysis. B. α/non-α ratio in the erythroblasts generated from gene edited HSPCs C. Representative globin chain HPLC chromatograms. D. Percentage of Annexin V in the erythroblasts generated from gene edited HSPCs. E. Representative flow cytometry image of Annexin V staining. F. Percentage of reticulocytes generated from gene edited HSPCs G. Representative flow cytometry plots of reticulocytes marked by CD235a^+^/Hoechst^-^.

Ineffective erythropoiesis, the classical phenotype of β-thalassemia eventuates due to increased reactive oxygen species (ROS) levels, apoptosis of erythroid progenitors, and reticulocyte maturation arrest(*3, 32*). Erythroblasts originated from PRR-βE1 gene edited group showed a modest decrease in ROS levels (Fig. S7D), a decrease in the proportion of apoptotic erythroblasts, stained by Annexin V (Fig. 6D-E) and importantly, up to 3-fold increase in reticulocyte generation (Fig. 6F-G) compared to control (AAVS1). All these findings suggest that PRR-βE1 gene editing functionally rescues erythropoiesis in β-thalassemia by robust activation of g-globin and silencing of b-globin (Fig. S9).

### PRR-βE1 gene editing reconfigures chromosome looping and alters globin expression in a HBBP1 dependent mechanism

Long-range chromatin interaction of Locus Control Region (LCR) and the promoters in the β-globin cluster regulate developmental stage specific expression of globin genes(*33*). To test the potential impact of PRR-βE1 gene editing in the configuration of β-globin cluster, we employed Circular chromosome conformation capture (4C) assay. An interaction between hypersensitive sites 1 (HS1) within LCR region and *HBG2* promoter was observed in HUDEP-2 control cells. However, this interaction was enhanced in HUDEP-2 clones harbouring PRR-βE1 biallelic deletion. Further, the interaction between other HS sites and HBG2 promoters was newly gained in PRR-βE1 deleted cells. (Fig. 7A). This data suggests that genomic proximity between LCR region and *HBG* gene increases upon PRR-bE1 deletions and thus, reactivates g-globin in edited cells.

**Figure 7.**
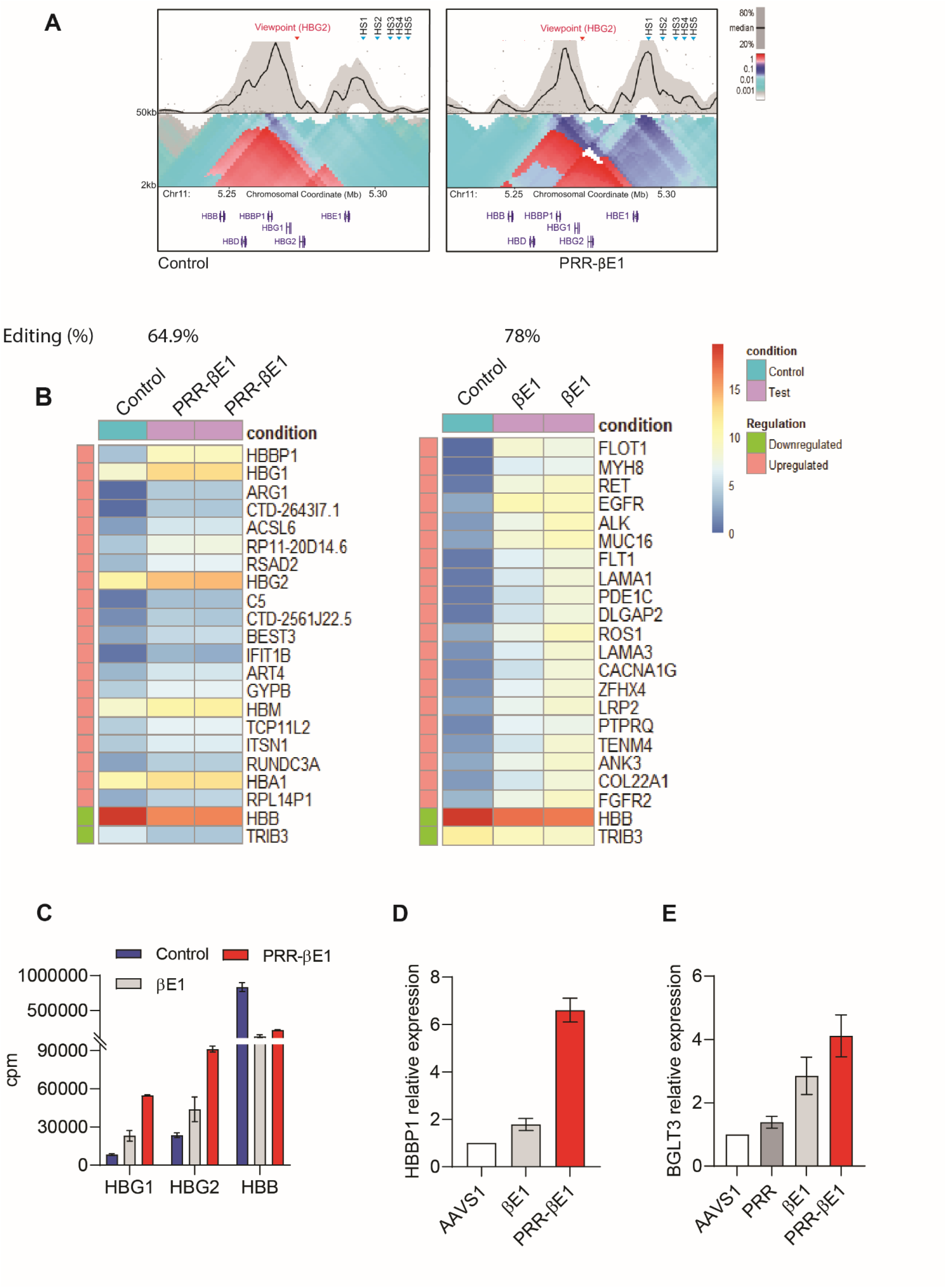
PRR-βE1 gene editing reconfigures chromosome looping and alters globin expression in a HBBP1 dependent mechanism. A. 4C analysis of single cell sorted control and PRR-βE1 gene edited HUDEP-2 clone using *HBG2* promoter as a viewpoint. B. Heat-map of erythroblasts derived from PRR-βE1 gene edited HSPC indicating the relative gene expression pattern of genes up and downregulated compared to control. C. Cluster per million (cpm) values for the globin transcripts obtained from RNA sequencing. D. Relative HBBP1 mRNA expression in erythroblasts derived from βE1 and PRR-βE1 gene edited HSPCs compared to AAVS1. The globin chain expression is normalised with β-actin. Donor =1, n = 4 E. Relative BGLT3 mRNA expression in erythroblasts derived from PRR, βE1, an PRR-βE1 gene edited HSPCs compared to AAVS1. The globin chain expression is normalised with β-actin. Donor =2, n = 4 Error bars represent mean ± SEM.

Thereafter, to understand the trans-acting factors involved in β-globin reactivation in the PRR-βE1 gene edited cells, transcriptome analysis was carried out using the erythroblasts generated *in vitro* from gene edited HSPCs. This analysis confirmed the overexpression of HBG1 and HBG2 with simultaneous downregulation of HBB (Figure 7B, C). The analysis also indicated the over expression of HBBP1, which was recently implicated in γ-globin activation(*34*). Real time PCR analysis confirmed the increased expression of HBBP1(Figure 7D). Earlier reports have demonstrated that another long non-coding RNA, BGLT-3, interacts with MED12 to promote transcriptional assembly at the γ-globin promoter(*35*). RT-PCR analysis also indicated that BGLT-3 transcripts were abundant in the edited cells (Figure 7E). All these data suggest that the γ-globin activation in PRR-βE1 gene edited cells occurs through altered chromatin looping mediated by promoter competition for LCR and the activation of long-noncoding RNAs HBBP1 and BGLT3.

To test whether our PRR-βE1 gene editing has resulted in any chromosomal abnormalities, we carried out array based KaryoStat analysis of edited and un-edited HSPCs. The edited HSPCs were expanded for 7-days to amplify any potential defect. The whole-genome coverage analysis with a resolution of >1 Mb indicated no loss or gain of chromosomal copy number in the PRR-βE1 gene edited HSPCs (Fig. S8C). This experiment suggests that the genome integrity is not disrupted upon PRR-βE1 gene editing. However, this experiment does not rule out the presence of on-target rearrangements in a small portion of cells.

## Discussion

Genetic reactivation of developmentally silenced fetal haemoglobin has gained considerable attention as a potential therapy for the broad spectrum of β-hemoglobinopathies. In this study, we have identified the PRR-βE1 sequence as a core HbF regulatory region present in all the deletional HPFH mutations. When present, PRR-βE1 disrupts defective β-globin production, and concurrently induces robust fetal haemoglobin production through HBBP1 dependent and LCR switching mechanism, thereby effectively reversing both SCD and β-thalassemia major phenotypes. In particular, we demonstrated that PRR-βE1 gene edited HSPCs possess long-term engraftment and repopulation fitness, underlining the potential of this approach for future clinical studies.

Among the naturally existing mutations that produce pancellular HbF, deletional HPFH mutations are highly prevalent and are shown to generate a high frequency of HbF^+ve^ red cells(*12*). Even a heterozygous deletion can result in HbF level of 65.6% with 8.9 g/dl of hemoglobin on co-inheritance with β-thalassemia(*19*).

Identifying the core region in HPFH deletions will enable us to recreate the HPFH phenotype by gene editing only the core region. The PRR region is conserved in δβ thalassemia but excised in HPFH deletions(*36*). However, deletion of the PRR site alone has not activated the fetal haemoglobin in our studies, consistent with earlier observations(*36*). Even a deletion of 10.5 kb spanning the PRR region to the region located before β-globin promoter had little effect on γ-globin production. On the other hand, disruption of βE1 alone induced γ-globin production. Shen et al showed that the improved γ-globin levels obtained by disrupting the HBB gene and its regulatory region is not sufficient to induce loss of β-globin(*37*). Our study provides strong evidence that simultaneous disruption of the PRR region and β-globin reactivates γ-globin robustly without generating any erythroid defects. While PRR region disruption ensures that we are not creating δβ-thalassemia like phenotype, the βE1 cut site associated γ-globin production makes PRR-βE1 more potent than the original Sicilian HPFH.

The ongoing clinical trial CLIMB SCD-121 showed that the SCD patient HSPCs gene edited for BCL11A erythroid-specific enhancer had gene editing efficiency up to 82.6% and HbF levels of 43.2% along with the presence of HbS tetramers up to 52.3% (*11*). This points out that irrespective of the gene editing efficiency and γ-globin activation efficacy, an intact β-globin regulatory region allows production of mutated β-globin chains at reduced levels. Similarly, in the BCL11A shRNA clinical trial, HbS constitutes up to 70% of hemoglobin tetramers(*10*). The PRR-βE1 editing strategy directly excises the promoter and coding regions of β-globin, resulting in a major reduction in the concentration of sickle haemoglobin which will prevent the sickling of red blood cells. The strategy will also be applicable for β-thalassemia where the intact β-globin promoter drives production of truncated β-globin chains (Fig. S9). To our knowledge, our study is the first to demonstrate the successful engraftment, multilineage differentiation, and serial repopulation of HSPCs harboring genomic deletion of > 5 kb.

## Materials and Methods

### Purification and culture of CD34^+ve^ HSPCs

The unused G-CSF mobilised peripheral blood collected for allogenic stem cell transplantation and Plerixafor mobilised peripheral blood from SCD, or β-thalassemia patients were collected from transplantation unit of Christian Medical College, Vellore with prior IRB approval. The PBMNCs were isolated using lymphoprep (STEMCELL Technologies) by density gradient centrifugation followed by isolation of CD34^+ve^ HSPCs using CD34 positive selection kit (STEMCELL Technologies) as per the manufacturer’s instructions. Prior to the nucleofection, the cells were pre-stimulated for 48-72hrs with StemSpan SFEM II (STEMCELL Technologies) containing 240ng/ml of SCF and Flt3-L, 80ng/ml of TPO, and 40ng/ml of IL6.

### Electroporation of RNP complex in HUDEP-2 and CD34^+ve^ HSPCs

SgRNAs were designed using CRISPR Design Tool (Synthego) and CRISPR-Cas9 guide RNA design checker (IDT), and the efficient gRNAs with least off-target sites were selected. List of gRNA used in the study are mentioned in sup. table 2. For nucleofection of HUDEP-2 cell lines, 100pmols of Cas9 (Takara) was incubated at room temperature for 10 minutes with 200pmols of sgRNA (Synthego). For dual sgRNA gene editing 100 pmols of Cas9 RNP with cut site A sgRNA and 100 pmols of Cas9 RNP with cut site B sgRNA were nucleofected (Lonza 4D nucleofector) with CA137 pulse code. For electroporation of CD34^+ve^ HSPCs, 50pmol of Cas9 RNP with sgRNA against PRR and 50pmol of Cas9 RNP with sgRNA against βE1 were used. 2×10^5^ cells were electroporated using P3 primary cell solution and supplement and were electroporated using Lonza 4D nucleofector with DZ100 pulse code.

### HUDEP-2 expansion and differentiation

The HUDEP-2 cells were cultured in StemSpan SFEM-II media containing SCF (50ng/ml), EPO (3U/ml), Dexamethasone (1µM), Doxycycline (1µg/ml), and Glutamine (1x) at 2×10^5^ cells/ml confluency with media change on alternative days. For erythroid differentiation, previously reported protocol with minor modifications was used(*38*). The cells were seeded at a density of 2×10^5^ cells/ml in IMDM GlutaMAX Supplement media containing 3% AB serum, 2% FBS, Insulin (10µg/ml), Heparin (3U/ml), EPO (3U/ml), Holotransferrin (200 µg/ml), SCF (100ng/ml), IL3 (10ng/ml) and Doxycycline (1µg/ml). On day 2, cells were seeded at a cell density of 3.5×10^5^ cells/ml. On day 4, the cells were seeded at a cell density of 5×10^5^ cells/ml in the media containing the above-mentioned cytokine except doxycycline. On day 6, the cells were seeded at a cell density of 1×10^6^ cells/ml in the media with all the components of day 4 media along with increased concentration of Holotransferrin (500 µg/ml). The cells were analysed for HbF^+ve^ cells, differentiation profile and globin chains using HPLC.

### Erythroid differentiation of CD34^+ve^ HSPCs

The protocol for erythroid differentiation from CD34^+ve^ HSPCs was adopted from the literature with minor modifications(*39*). The three-phase erythroid differentiation protocol involves culturing the CD34^+ve^ cells at a seeding density of 5×10^4^ cells/ml in phase I from day 0 – day 8 with a media change on day 4. The phase I media is prepared using IMDM GlutaMAX Supplement media containing 5% AB serum, Insulin (20µg/ml), Heparin (2U/ml), EPO (3U/ml), Holotransferrin (330 µg/ml), SCF (100ng/ml), IL3 (50ng/ml) and Hydrocortisone (1µg/ml). In phase II, the cells were seeded at a density of 2×10^5^ cells from day 8 – day 12 in media containing all the components of phase I except Hydrocortisone and IL3. In phase III, the cells were seeded at a density of 5×10^5^ cells from day 12 – day 20 in media containing all the components of phase II except SCF with a media change on day 14. On day 20, the cells were collected for F^+ve^ cells analysis, differentiation marker analysis and for haemoglobin and globin chain HPLC.

### Flow cytometry

For HbF^+ve^ cell analysis, 1×10^5^ erythroid differentiated cells were briefly washed with PBS and fixed with 0.05% glutaraldehyde for 10 minutes and permeabilized with 0.1% Triton-X-100 for 5 minutes. The cells were stained with anti-HbF APC antibody (dilution 1:50) and was acquired and analysed using Cytoflex LX Flow Cytometer (Beckmann Coulter) or AriaIII flow cytometer (BD Biosciences) and analysed using FlowJo (BD Biosciences). For erythroid differentiation analysis, 1×10^5^ cells from the terminal day of erythroid differentiation were stained for erythroid differentiation markers anti-CD71-FITC (dilution 1:33), anti-CD235a PE-Cy7 (dilution 1:50) and Hoechst 33342 (dilution 1:1000). After 20 minutes of incubation in dark, the cells were washed with PBS followed by the analysis using Cytoflex LX Flow Cytometer (Beckmann Coulter) or AriaIII flow cytometer (BD Biosciences).

### *In vivo* engraftment analysis

All the *in vivo* experiments in mice models (NSG and NBSGW) were conducted with approval form IAEC of Christian Medical College, Vellore, India. Both the NSG and NBSGW were bred in in-house animal facility. For conditioning, 7-9 weeks old female NSG mice were irradiated sub-lethally with 250cGy 8-12hrs prior to infusion and NBSGW mice were conditioned with busulfan at a concentration of 12.5mg/kg of body weight, 48hrs prior to the infusion. CD34^+^ HSPCs were pre-stimulated for 36hrs – 40hrs with culture media containing appropriate cytokines and RUS cocktail(*40*). 5×10^5^ -6×10^5^ cells of control edited and PRR-βE1 edited were infused in NSG and NBSGW mice, immediately post electroporation. 16-18 weeks post infusion, the mice were euthanised and peripheral blood, bone marrow and spleen were collected. After RBC lysis buffer incubation, the harvested cells were incubated with mouse Fc block and stained with hCD45 and mCD45 antibody. The % of engraftment is calculated using the formula (% hCD45/% hCD45+% mCD45) x 100. In addition, the multilineage markers including CD19, CD3, CD33, CD13, and CD235a in bone marrow hCD45+ cells were also analysed. For *ex vivo* erythroid differentiation, 3×10^6^ cells were harvested from mouse bone marrow, seeded in erythroid differentiation media and at the end of phase III of differentiation, the % of F+ve cells, the differentiation profile and globin chains were analysed. For *in-vivo* HbF^+ve^ cell analysis in NBSGW, 1×10^6^ bone marrow cells were stained with 10µL of CD235a antibody and sorted based on the presence of immunophenotypic marker CD235a and was followed by F^+ve^ cell analysis. For secondary infusion, 3×10^6^ cells from the pooled fraction harvested from primary recipient bone marrow were infused to secondary recipients at a time point of 8-12hrs post irradiation. After 8 weeks, the mice were euthanised and the harvested cells were stained with hCD45 and mCD45 antibody for calculating the % of engraftment.

### Quantitative Real-Time PCR Analysis

3×10^6^ cells from the day 8 of CD34^+^ HSPC and day 6 of HUDEP-2 erythroid differentiation were used for total RNA using RNeasy Mini Kit (Qiagen). For reverse transcription using Primescript RT reagent kit (Takara Bio Inc.), 1 μg of extracted RNA was used according to manufacturer’s instruction. For quantitative PCR, the SYBR Premix Ex Taq II (Takara Bio) was used for quantifying the specific transcripts and analyzed with QuantStudio 6 Flex (Life Technologies). Primers used in qPCR analysis are mentioned in sup. table 5.

### Colony formation assay

48hrs post electroporation, 5×10^2^ HSPCs were seeded in 1.5ml of Methocult Optimum (STEMCELL Technologies) and after 14 days, the colonies were scored based on the morphology as CFU-GM, CFU-GEMM, BFU-E, and CFU-E.

### Digital droplet PCR (ddPCR)

The frequency of PRR-βE1 genomic deletions were quantified using EvaGreen based ddPCR assay. The reaction mixture includes 20ng of genomic DNA, 1x QX200 ddPCR EvaGreen supermix and 100nM primers for 20µL reaction. For absolute measure of deletions, we designed primers that amplify the sequences flanking the cut sites after targeted deletion. Control primers amplifying embryonic globin gene were used as loading control (EG). The percentage of deletion was calculated using the formula.

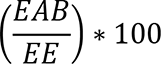

Where,

**EAB** – DNA copies/µl from primers flanking the cut sites of edited samples.

**EG** - DNA copies/µL from primers amplifying embryonic globin gene of edited samples.

The second approach involves the quantification of the individual cut sites of the deletion, normalised with the read outs from the unedited control samples.

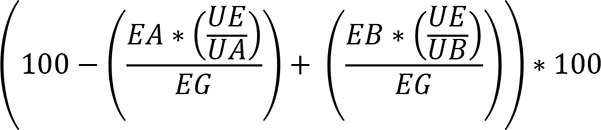

Where,

**EA** – DNA copies/µL from primers flanking the cut site A of edited samples.

**EB** – DNA copies/µL from primers flanking the cut site B of edited samples.

**EG** - DNA copies/µL from primers amplifying embryonic globin gene of edited samples.

**UA** – DNA copies/µL from primers flanking the cut site A of unedited samples.

**UB** – DNA copies/µL from primers flanking the cut site B of unedited samples.

**UE** - DNA copies/µL from primers amplifying embryonic globin gene of unedited samples.

Cut site A indicates PRR region and Cut site B indicates βE1 region. Primers used in ddPCR analysis are mentioned in sup. table 4.

### Haemoglobin and globin chain analysis using High performance liquid chromatography

The gene edited HUDEP-2 cell lines and CD34^+ve^ HSPCs were collected on day 8 and day 20 of erythroid differentiation, respectively. The cells were sonicated for 60 seconds with 50% AMP in ice using ultrasonicator (Vibra-Cell) and centrifuged at 14000 rpm for 5 minutes at 4°C. For haemoglobin HPLC, the protein lysate was analysed for haemoglobin tetramer using G8 HPLC Analyzer (Tosoh). The globin chain analysis was performed using HPLC equipment with UV detector (Shimadzu) and the analysis was performed using LC SolutionsTM software (Shimadzu) using previously reported method(*41*). Aeris Widepore 3.6 lm XB-C18 25cm 4.6mm column behind a Security Guard UHPLC Widepore C18 4.6mm guard column (PhenomenexTM) is used for chromatographic separation of the analytes. HPLC conditions include 0.1% trifluoroacetic acid (TFA), pH 3.0 (solvent A), mobile phase - 0.1% TFA in acetonitrile (solvent B) with gradient elution at a flow rate of 1.0 ml/min and column temperature maintained at 70° C with runtime around 8 minutes and UV detection range of 190nm was set for globin chain detection.

### Western Blot Analysis

Approximately 6×10^6^ erythroblasts were collected on day 8 of erythroid differentiation. The lysates were prepared sonicating the cell pellets resuspended in RIPA buffer supplemented 1x protease and phosphatase inhibitor cocktail at 50% amplitude for 60 seconds as 30 second intervals in sonicator. 20µg of protein lysates resuspended in 1x *in house* prepared Lamelli buffer were loaded to the wells of SDS PAGE. The western blots were performed using the primary antibodies, anti-hemoglobin α (1:1000 dilution), anti-hemoglobin β (1:1000 dilution), anti-hemoglobin γ (1:1000 dilution) and anti-actin (1:5000 dilution) along with anti-mouse IgG HRP secondary antibodies. Densitometric analysis of the globin bands were performed by normalising with β-actin.

### Transcriptome analysis

Total RNA was extracted using Qiagen RNA isolation kit, quantified using Qubit RNA Assay HS, purity checked using QIAxpert and RNA integrity was assessed on TapeStation using RNA HS ScreenTapes (Agilent, Cat# 5067-5579). NEB Ultra II Directional RNA-Seq Library Prep kit protocol was used to prepare libraries for total RNA sequencing. Prepared libraries were quantified using Qubit High Sensitivity Assay (Invitrogen, Cat# Q32852). A cluster flow cell is loaded on Illumina HiSeq 4000 instrument to generate 60M, 100bp paired end reads. Read Counts from mapped reads were obtained using Feature Counts. Differential expression analysis was performed using DESEQ2. Gene set enrichment analysis was by GSEA software from the Broad Institute. A ranked list of differentially expressed genes from RNAseq data was loaded into GSEA and tested against a list of genes documented from published reports. Heat map for differentially regulated genes were generated using Morpheus (broad institute).

### KaryoStat assay

Gene edited HSPCs were collected for genomic DNA isolation using PureLink Genomic DNA Mini Kit (catalog # K182000) and quantified using Qubit dsDNA assay. After digestion of 250ng of genomic DNA using Nsp I restriction enzyme, the DNA were ligated with adapter and amplified. The DNA were fragmented followed by labelling with biotin and the labelled DNA was hybridised onto GeneChip arrays. GeneChip Fluidics Station 450 were used for washing and staining of Chips simultaneously scanned using GeneChip Scanner 3000 7 G. Data were analyzed using ChAS 3.2. The raw data were processed using Genotyping Console v4.0 and Chromosome Analysis Suite 3.2 with NetAffx na33.1 (UCSC GRCh37/hg19), and the output data were interpreted with the UCSC Genome Browser (https://genome.ucsc.edu/; GRCh37/hg19 assembly).

### 4C analysis

4C was performed as per the protocol described in the literature with minor variations(*42*). Cells (HUDEP-2 control and PRR-βE1 clone) were fixed with fresh formaldehyde (1.5%) and quenched with glycine (125 mM) followed by washes with ice-cold PBS (2×) and pelleted and stored at −80 °C. Lysis buffer [Tris-Cl pH 8.0 (10 mM), NaCl (10 mM), NP-40 (0.2%), PIC (1×)] was added to the pellets and were homogenized by Dounce homogenizer (15 stroked with pestle A followed by pestle B). The 3C digestion was performed with Csp6I (10 units, Thermofisher #ER0211) and ligation was performed by the T4 DNA ligase in 7.61 ml ligation mix (745 μl 10% Triton X-100, 745 μl 10x ligation buffer (500 mM Tris-HCl pH7.5, 100 mM MgCl2, 100 mM DTT), 80 μl 10 mg/ml BSA, 80 μl 100 mM ATP and 5.96 ml water). The ligated samples were de-crosslinked overnight then purified by PCI purification and subjected to ethanol precipitation and the pellet was eluted in TE (pH 8.0) to obtain the 3C library. The second 4C digestion was performed by DpnII (50 units, NEB) and the samples were ligated, purified and precipitated similar to the 3C library to obtain the 4C library. The 4C library was subjected to RNAseA treatment and purified by the QIAquick PCR purification kit. The concentration of the library was then measured by Nanodrop and subjected to PCRs using the oligos for the respective viewpoints. The oligos used for the HBG2 viewpoint are mentioned in the (sup. table 6). The samples were PCR purified and subjected to next-generation sequencing with Illumina HiSeq2500 using 50 bp single-end reads. Data analysis was performed using 4Cseqpipe (https://github.com/changegene/4Cseqpipe) using default parameters.

## Acknowledgments

The authors thank the help of Dr. Sowmya Pattabhi in designing the ddPCR strategy, staffs of flow cytometry, animal, and core facilities for the support.

## Author contributions

Conceptualization: ST, AS, SV, SM and KM.

Experiment execution and analysis: VV, AC, PB, MA, KW, BS, SS, KVK, AJ, SR, AP, YN, RP.

Technical Supervision: ST, AS, DN, SV, PB, SM.

Manuscript –review & editing: VV, ST, AS, SV, SM and KM.

Funding Acquisition; ST.

## Data and materials availability

All data are available in the main text or the supplementary materials. Additional data related to this paper may be requested from corresponding author.

## Supplementary figures

**Supplementary figure 1:**
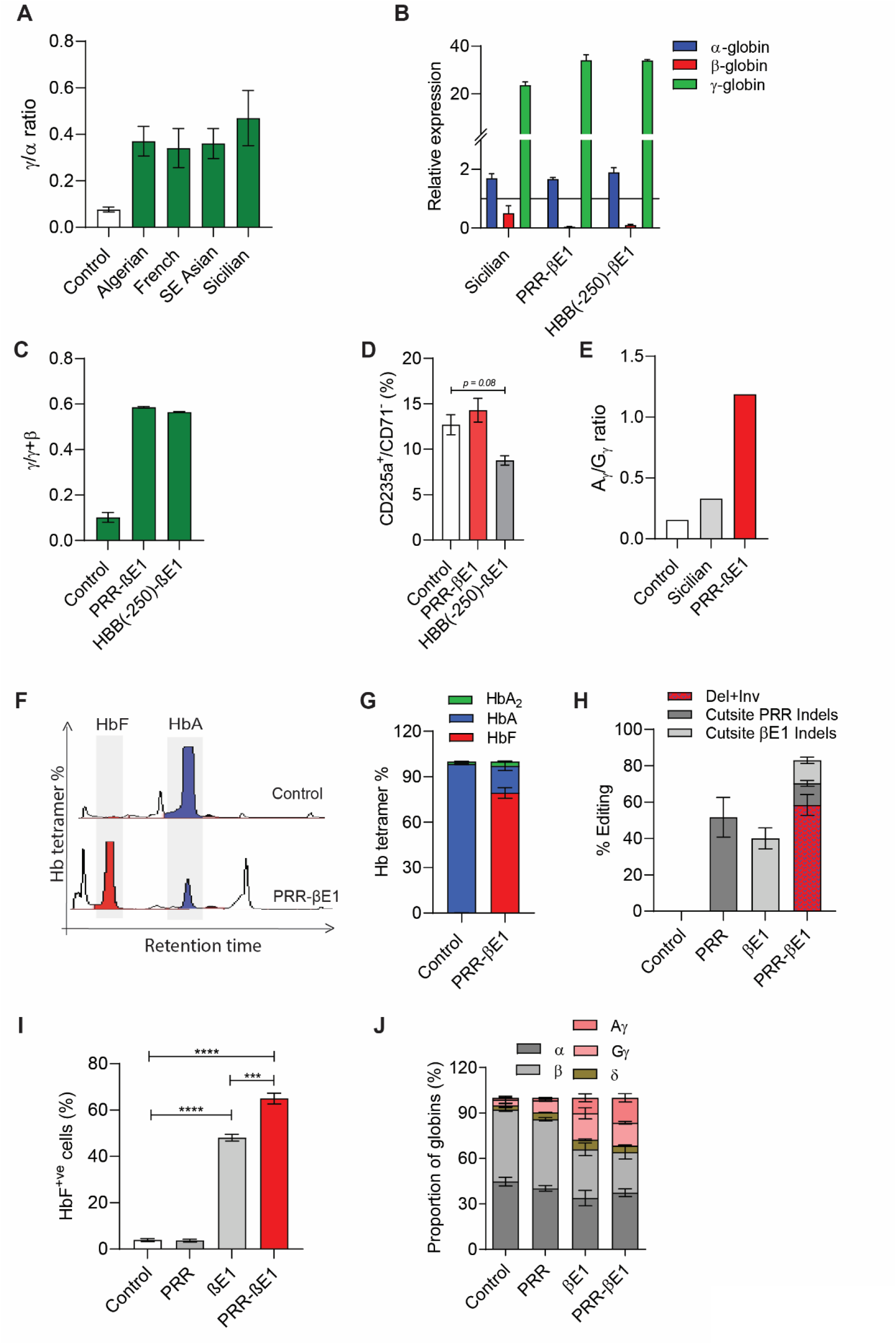
Genomic deletion encompassing PRR and exon1 of β-globin is sufficient for deletional HPFH phenotype. A. Ratio of γ/α and in HUDEP-2 cell lines gene edited for naturally occurring HPFH deletions and differentiated into erythroblasts. Globin chain activation was measured by HPLC chain analysis. n = 3. B. Relative globin mRNA expression in Sicilian, PRR-βE1, and HBB(-250)-βE1 gene edited HUDEP-2 cells compared to un-edited control. The globin chain expression is normalised with β-actin. n = 2. C. Ratio of γ/γ+β in HUDEP-2 cell lines gene edited for PRR-βE1, and HBB(-250)-βE1 and differentiated into erythroblasts. Globin chain activation was measured by HPLC chain analysis. n = 2. D. Percentage of CD235a^+^/CD71^-^ cells generated on erythroid differentiation of HUDEP-2 cell lines gene edited for PRR-βE1, and HBB(-250)-βE1. The cells were analysed for erythroid differentiation on day 8. n = 2. E. Ratio of Aγ/Gγ in HUDEP-2 cell lines gene edited for naturally occurring HPFH deletions and differentiated into erythroblasts. Globin chain activation was measured by HPLC chain analysis. n = 1. F. Representative hemoglobin variant HPLC chromatograms showing HbF and HbA tetramers in the control, PRR-βE1 gene edited and erythroid differentiated HUDEP-2 cells. G. Proportion of hemoglobin tetramer in control, PRR-βE1 gene edited and erythroid differentiated HUDEP-2 cells. n =3. H. Percentage of gene editing in PRR, βE1, and PRR-βE1 gene edited HUDEP-2 cell lines. Indels measured by ICE analysis of sanger reads. Deletion (red checker box) in PRR-βE1 quantified by ddPCR. The PRR-βE1 edited cells had deletion, indels at PRR region (cut site A indels) and βE1(cut site B indels). n =3. I. Percentage of HbF^+ve^ cells in PRR, βE1, and PRR-βE1 gene edited HUDEP-2 cell lines and differentiated into erythroblasts and analyzed HbF^+ve^ cells on day 8 of erythroid differentiation. n = 4. J. Proportion of globin chains as measured by chain HPLC in control and PRR-βE1 gene edited and erythroid differentiated HUDEP-2 cells. n =4 Error bars represent mean ± SEM, ^∗∗∗∗^p ≤ 0.0001 (Multiple t test).

**Supplementary figure 2:**
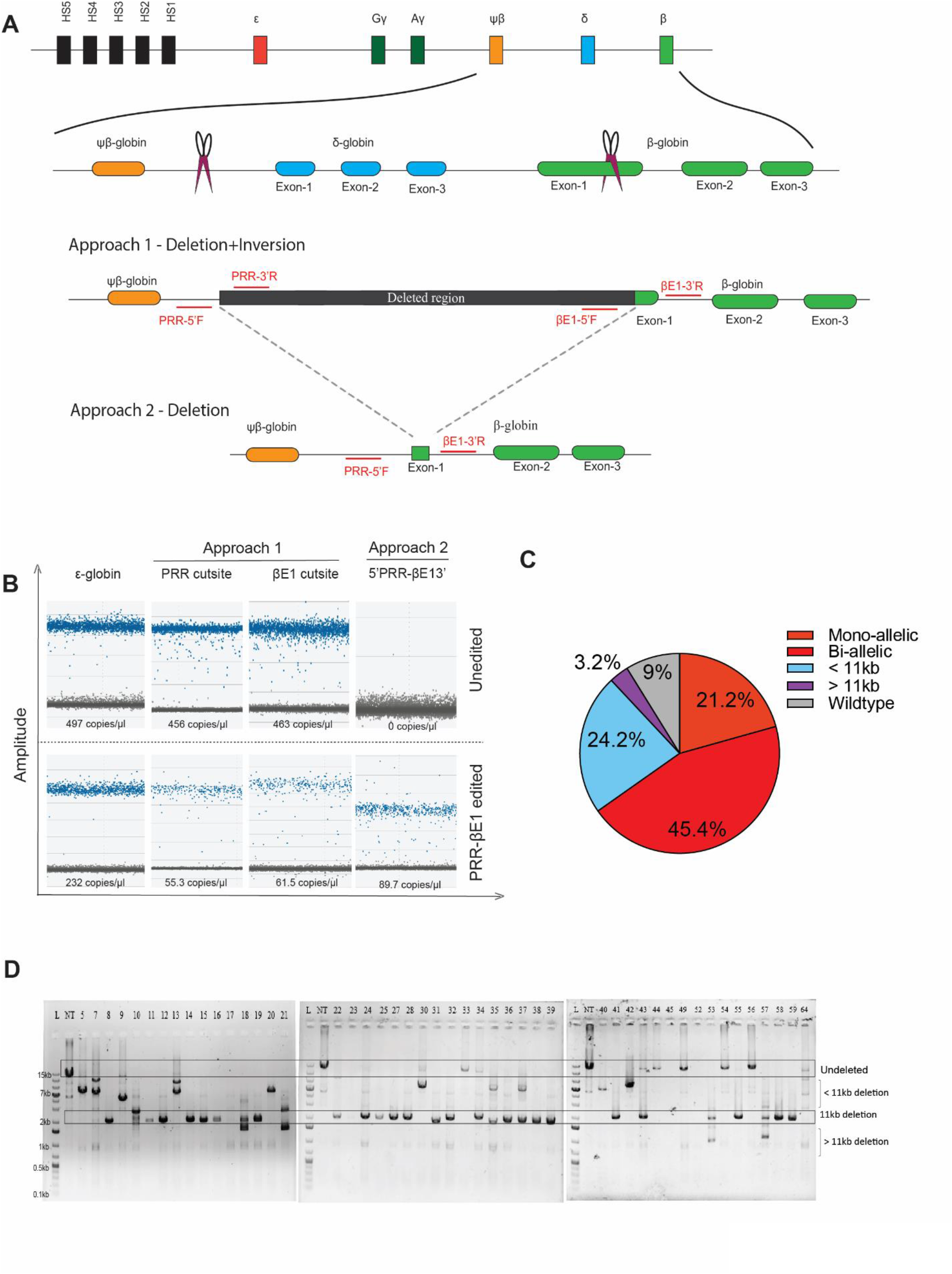
Quantification of PRR-βE1 genomic deletion. A. Diagrammatic representation of ddPCR approaches employed for quantifying the PRR-βE1 deletion events. In approach 1, EvaGreen dye based ddPCR assay was employed to quantify the loss of genomic regions spanning the cut site A (PRR) and cut site B (βE1). The percentage of deletion and inversion is calculated by comparing the number of positive droplets generated for control region (ε-globin) of the gene edited samples to the number of positive droplets produced on targeting the individual cut sites. The difference in the positive droplets is further normalised with the positive droplets of the cut sites of un-edited sample. In approach 2, the ddPCR assay was employed to quantify the ligated ends after excision of targeted 11kb deletion. The percentage of deletion is calculated by comparing the number of positive droplets generated for control region (ε-globin) of the gene edited sample to the number of positive droplets produced on targeting the ligated region. B. Representative ddPCR amplitude plot depicting the positive droplets and copy numbers of control region (ε-globin) compared to loss of positive droplets in PRR cut site and βE1 cut site due to PRR-βE1 deletion (approach 1) and amplification of ligated ends (approach 2). C. Frequency of deletions in single cell sorted HUDEP-2 cells gene edited for PRR-βE1 deletion as measured by gap PCR. D. Representative Gap PCR image of single cell clones of PRR-βE1 gene edited HUDEP-2 cells. Unedited samples shows amplicon size of 14.2kb and on PRR-βE1 deletion, primer amplifies the ligated regions spanning the cut site resulting in 2.1kb amplicon.

**Supplementary figure 3:**
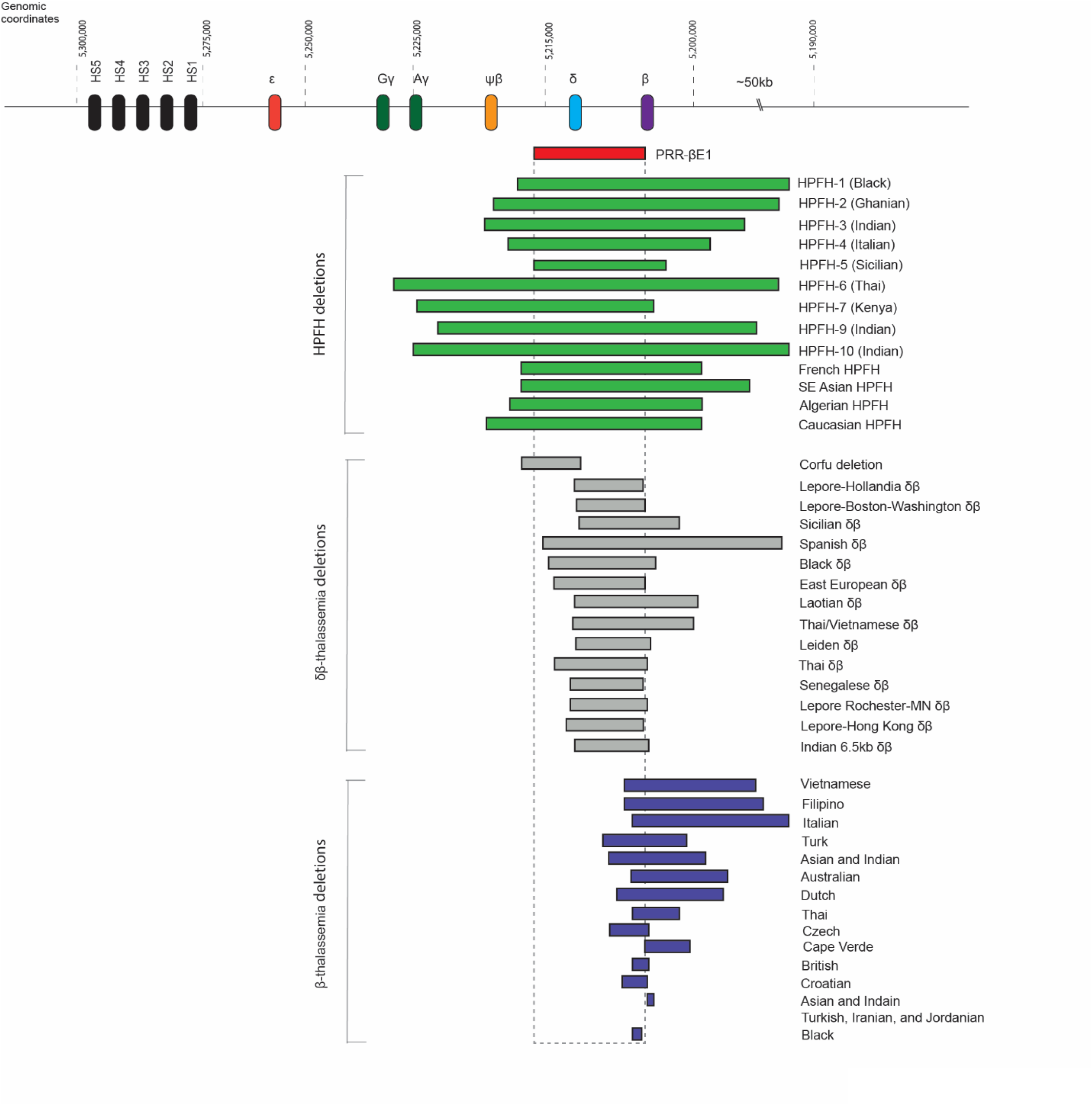
Genomic mapping of deletions in deletional HPFH mutations, δβ-thalassemia, and β-thalassemia. Comparison of genomic deletions in naturally occurring deletional HPFH mutations (green), δβ-thalassemia (grey), and β-thalassemia (blue) with PRR-βE1 deletion (red) indicates that the 11kb PRR-βE1 deletion remains as the key core region (highlighted in box) in all the HPFH deletions distinguishing the pan cellular and hetero cellular HbF producing deletions.

**Supplementary figure 4:**
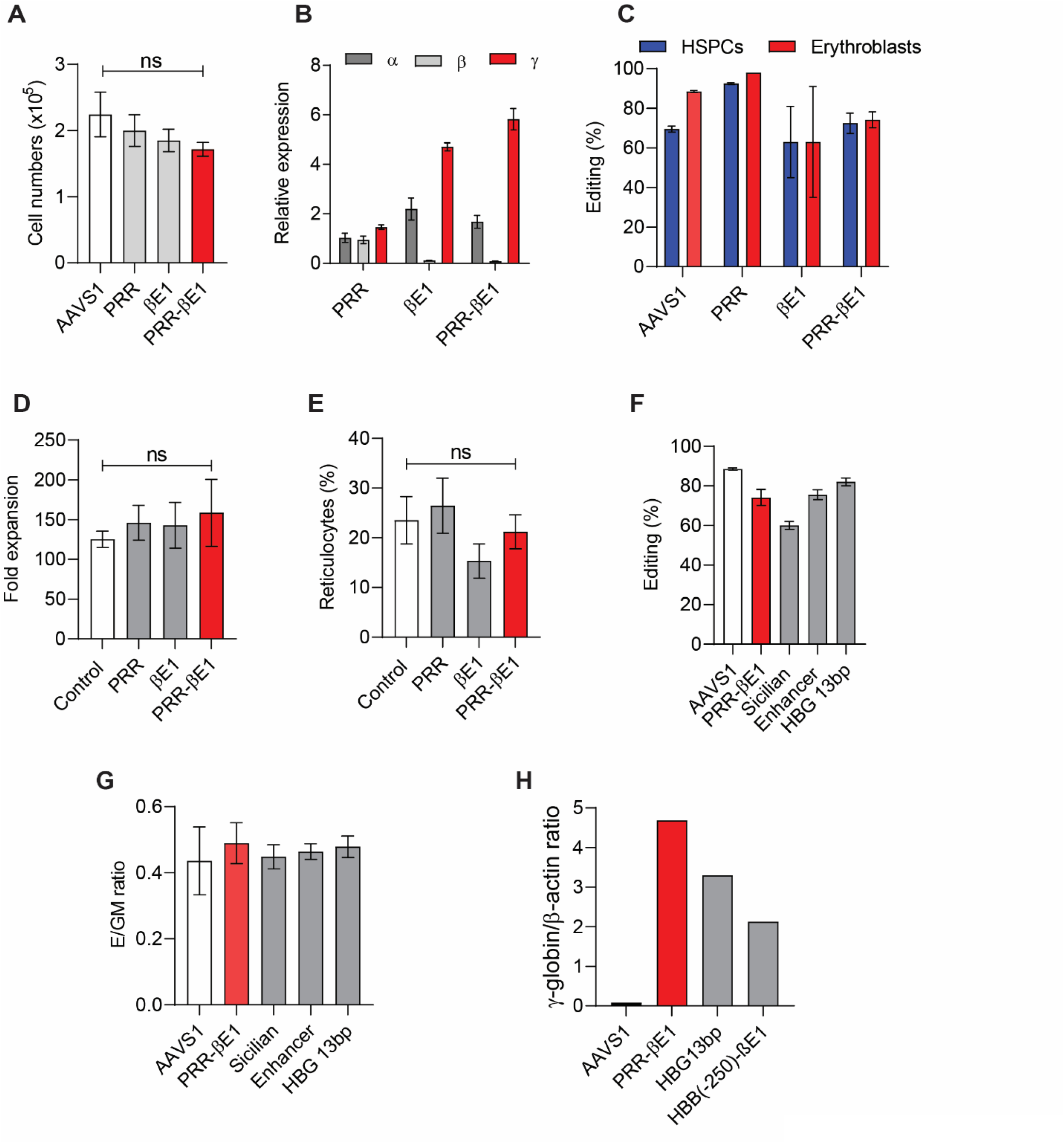
Intact erythropoiesis from PRR-βE1 gene edited HSPCs. A. Live cell numbers of AAVS1, PRR, βE1, and PRR-βE1 edited HSPCs counted 72hrs post electroporation using trypan blue. Donor = 4, n = 8. B. Relative globin mRNA expression on erythroid differentiation of PRR, βE1, and PRR-βE1gene edited HSPCs compared to AAVS1. The globin chain expression was normalised with β-actin. Donor =1, n = 4. C. Percentage of PRR-βE1 deletion in HSPCs and in erythroblasts. Deletion was measured by ddPCR. Donor = 1, n =2. D. Fold expansion of erythroblasts on day 12 of erythroid differentiation. Donor = 2, n =4. E. Percentage of reticulocytes generated on erythroid differentiation of HSCPs gene edited for PRR, βE1 and PRR-βE1. Donor =4, n = 7. F. Percentage of gene manipulation as measured by ddPCR for quantifying deletions in PRR-βE1 and Sicilian HPFH. Indel analysis of AAVS1, BCL11A enhancer, and HBG 13bp by ICE analysis. Donor =1, n =2. G. Erythroid to granulocyte-monocyte (GM) ratio of HSPCs. HSPCs gene edited for PRR-βE1, Sicilian HPFH, BCL11a enhancer and HBG 13bp promoter was analysed for CFU potential. Donor = 2, n = 4. H. Densitometric analysis of γ-globin/β-actin ratio for HSPCs derived erythroblasts gene edited for AAVS1, PRR-βE1, HBG13bp, and HBB(-250)-βE1. Donor = 1, n = 1. Error bars represent mean ± SEM, ns; non-significant (Multiple t test).

**Supplementary figure 5:**
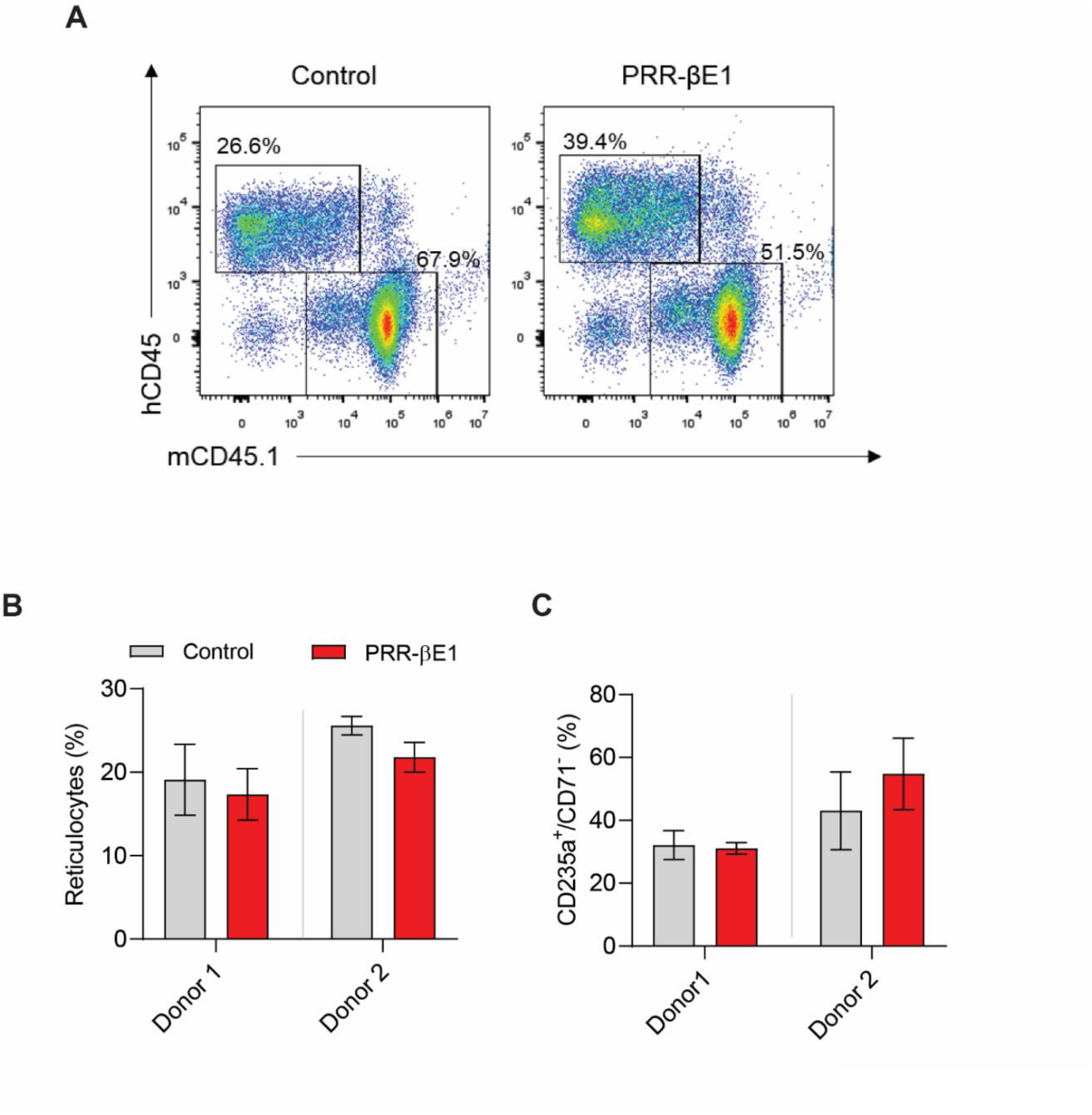
Engraftment characteristics of PRR-βE1 gene edited HSPCs. A. Representative flow cytometry image of hCD45 and mCD45.1 cells in mouse bone marrow of NSG mice. B. Percentage of reticulocytes generated on *in vitro* erythroid differentiation of bone marrow mononuclear cells in NSG mice infused with Control and PRR-βE1 gene edited HSPCs. Donor = 2, n = 4 C. Percentage of CD235a^+^/CD71^-^ cells generated on *in vitro* erythroid differentiation of bone marrow mononuclear cells in NSG mice infused with Control and PRR-βE1 gene edited HSPCs. Donor = 2, n = 4

**Supplementary figure 6:**
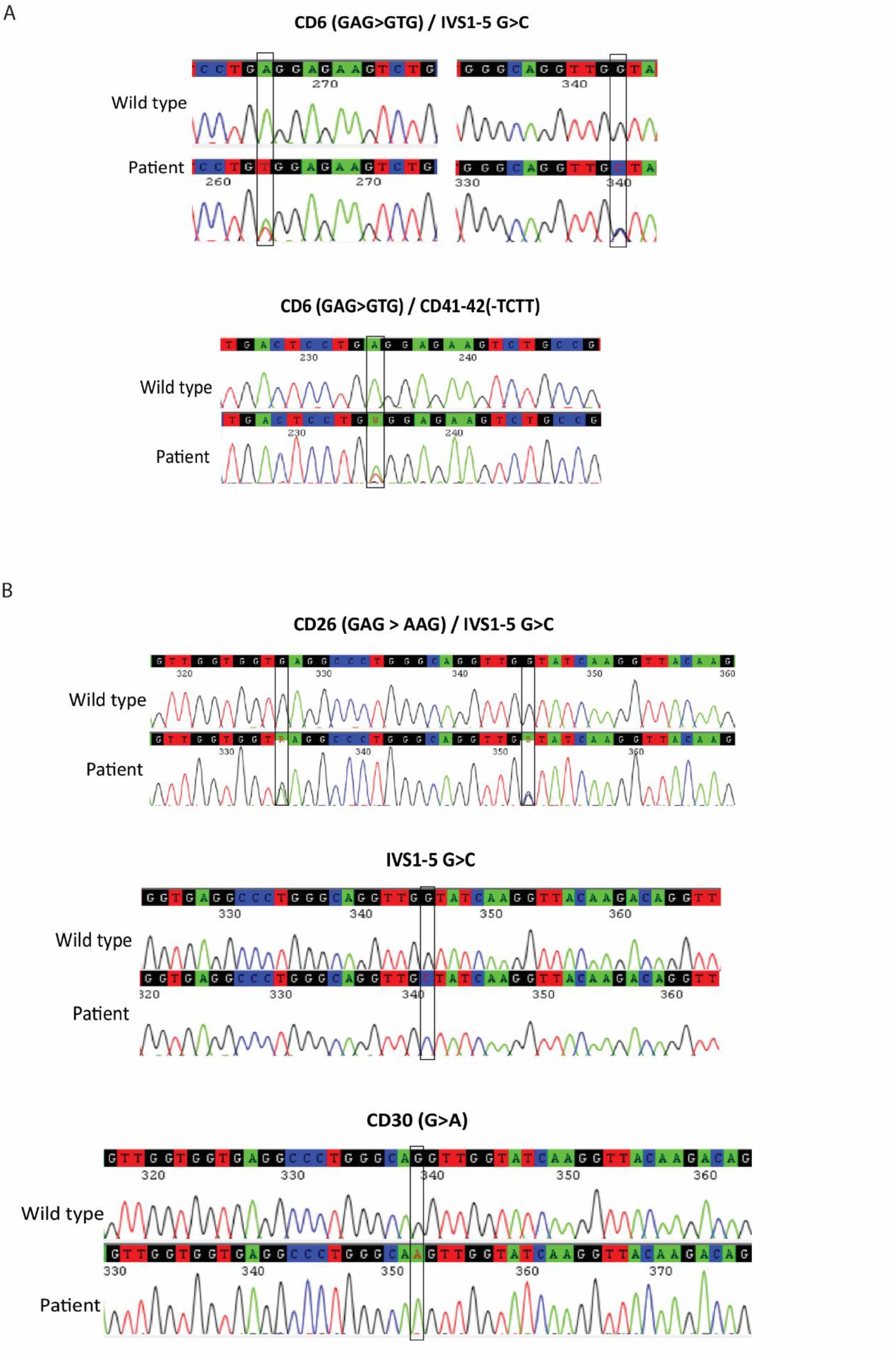
Genotype characterisation of patient HSPCs. A. Genotype of SCD patient HSPCs identified by Sanger sequencing. B. Genotype of β-thalassemia patient HSPCs identified by Sanger sequencing.

**Supplementary figure 7:**
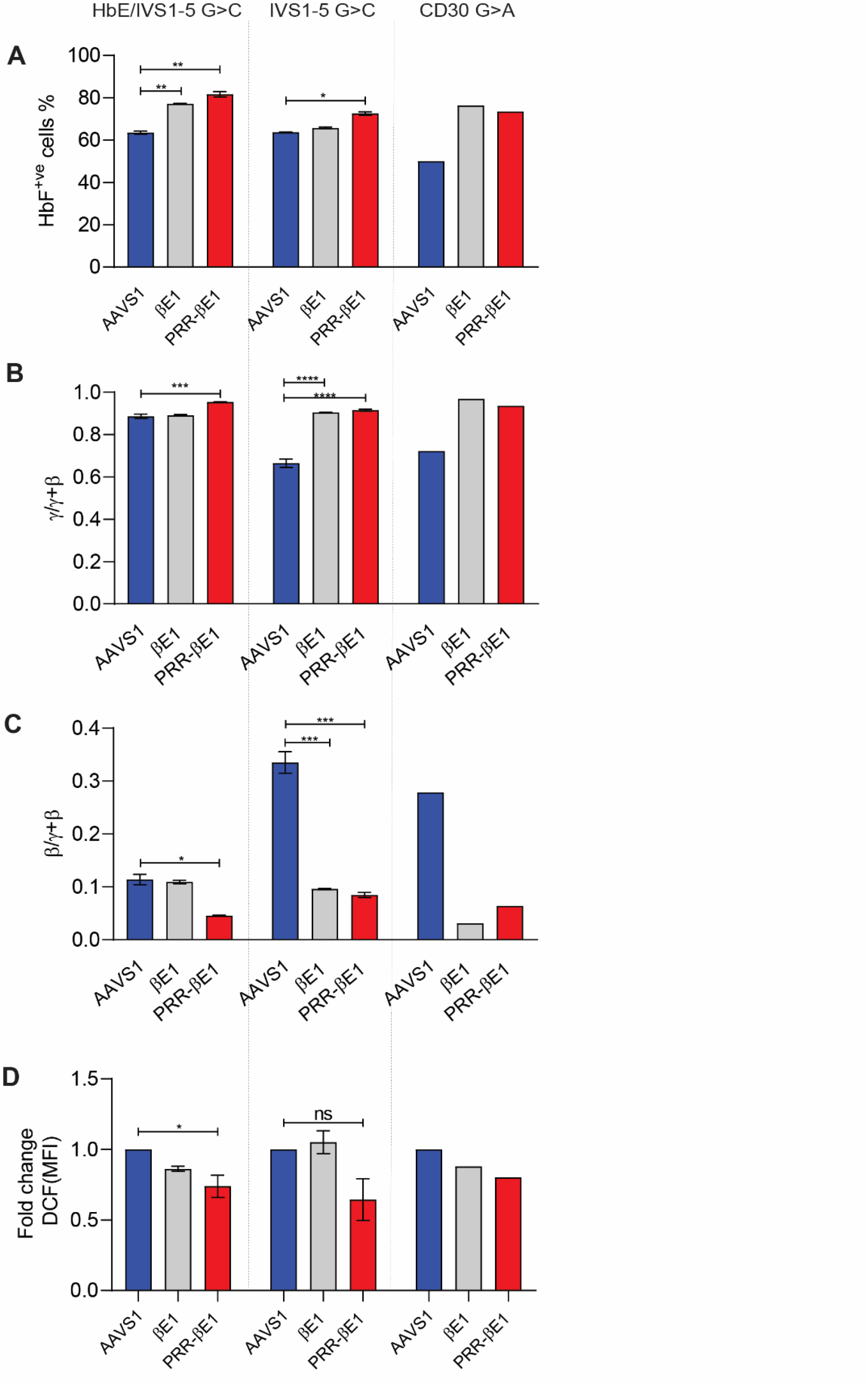
β-hemoglobinopathies reversal characteristics of PRR-βE1 gene edited HSPCs. A. Percentage of HbF^+ve^ cells in erythroblasts derived from gene edited β-thalassemia patient HSPCs. Donor = 3, n = 6 B. γ**/**γ**+**β ratio in erythroid differentiated G-CSF mobilised patient HSPCs gene edited for PRR, βE1, and PRR-βE1. Donor = 3, n = 6. C. β**/**γ**+**β ratio in erythroid differentiated G-CSF mobilised patient HSPCs gene edited for PRR, βE1, and PRR-βE1. Donor = 3, n = 6. D. Fold change in the DCF mean fluorescence intensity in erythroblasts generated from gene edited β-thalassemia patient HSPCs. Donor = 3, n = 6 Error bars represent mean ± SEM, ns; non-significant. ^∗^p ≤ 0.05, ^∗∗^p ≤ 0.01 (Multiple t test).

**Supplementary figure 8:**
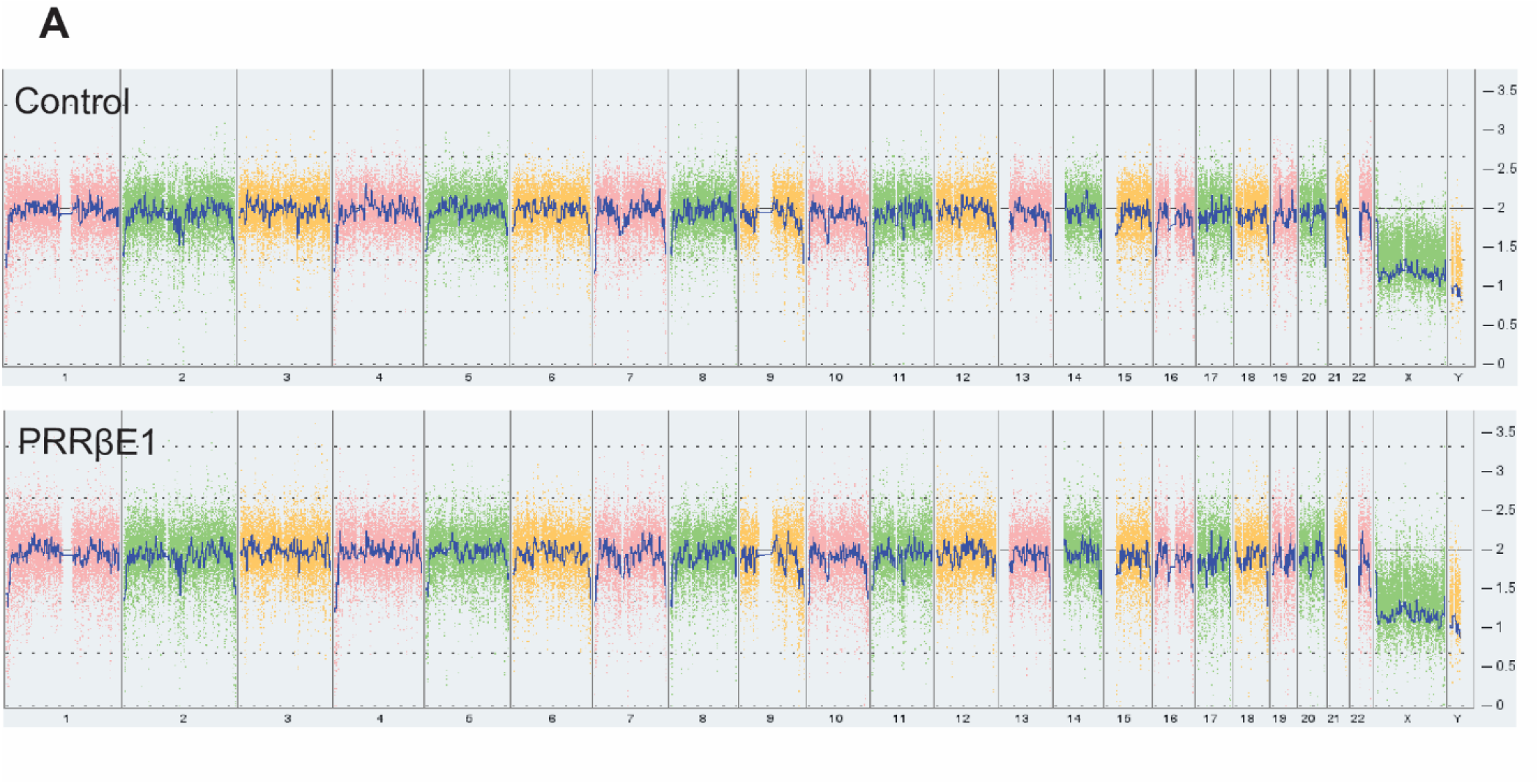
Genome integrity characterisation of of PRR-βE1 gene edited HSPCs. A. KaryoStat analysis of healthy donor HSPCs gene edited for PRR-βE1 deletion. Donor = 1, n = 2.

**Supplementary figure 9:**
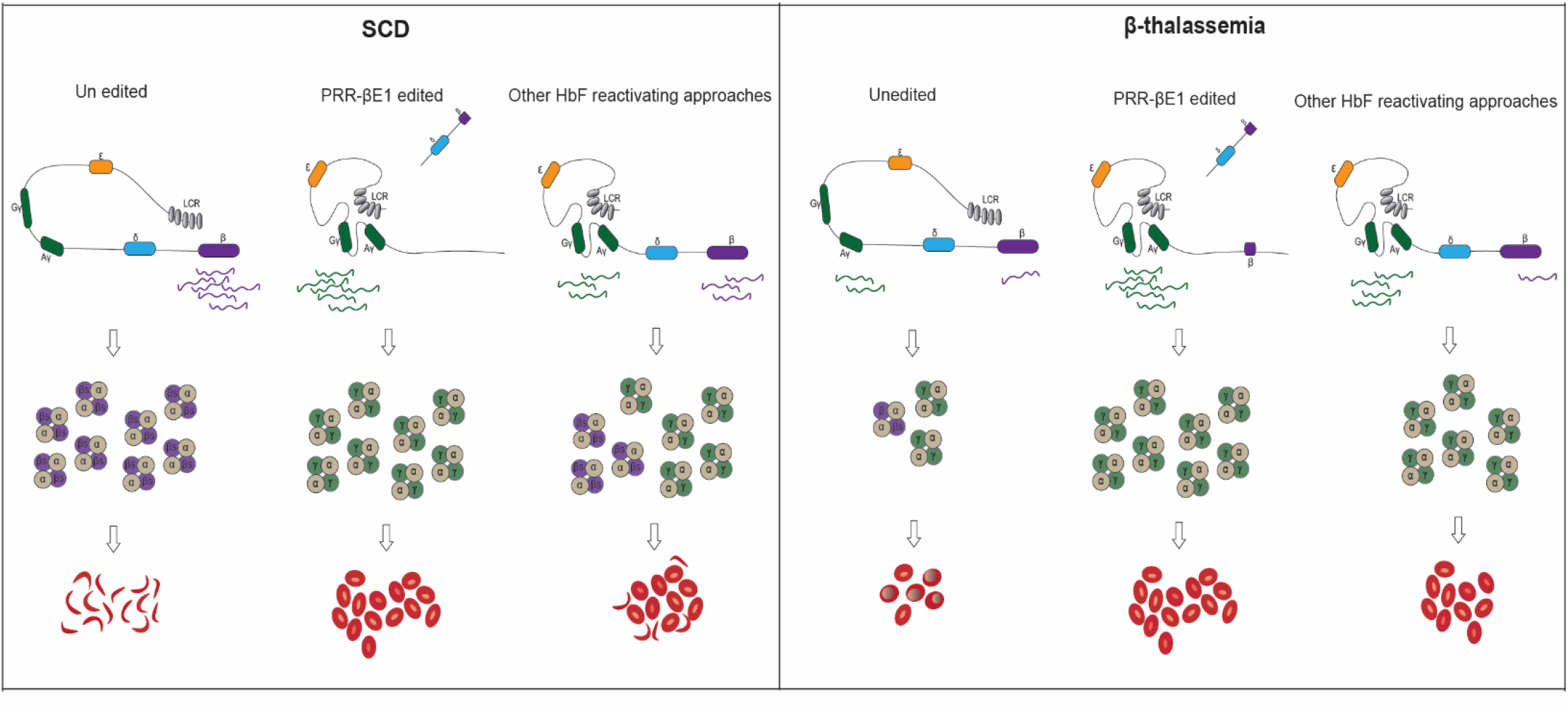
Amelioration of SCD and β-thalassemia phenotype on PRR-βE1 deletion. Graphical representation on amelioration of SCD and β-thalassemia phenotype on PRR-βE1 deletion

## Supplementary tables

**Table – S1.**

| ANTIBODY/DYE | COMPANY | CLONE | CAT NO |
| --- | --- | --- | --- |
| CD45.1 PerCP Cy5.5 | BD | A20 | 560580 |
| CD3 PECY 7 | BD | SP34-2 | 557749 |
| CD13 PE | BD | WM15 | 555394 |
| CD71 FITC | BD | M-A712 | 555536 |
| CD235a PE Cy-7 | BD | GA-R2(HIR2) | 563666 |
| CD45 APC | BD | HI30 | 555485 |
| CD34 PE | BD | 8G12 | 348057 |
| CD19 PerCP | BD | - | 340421 |
| Hoechst 33342 | Thermo Scientific |  | 62249 |

List of antibodies used in the study

**Table – S2.**
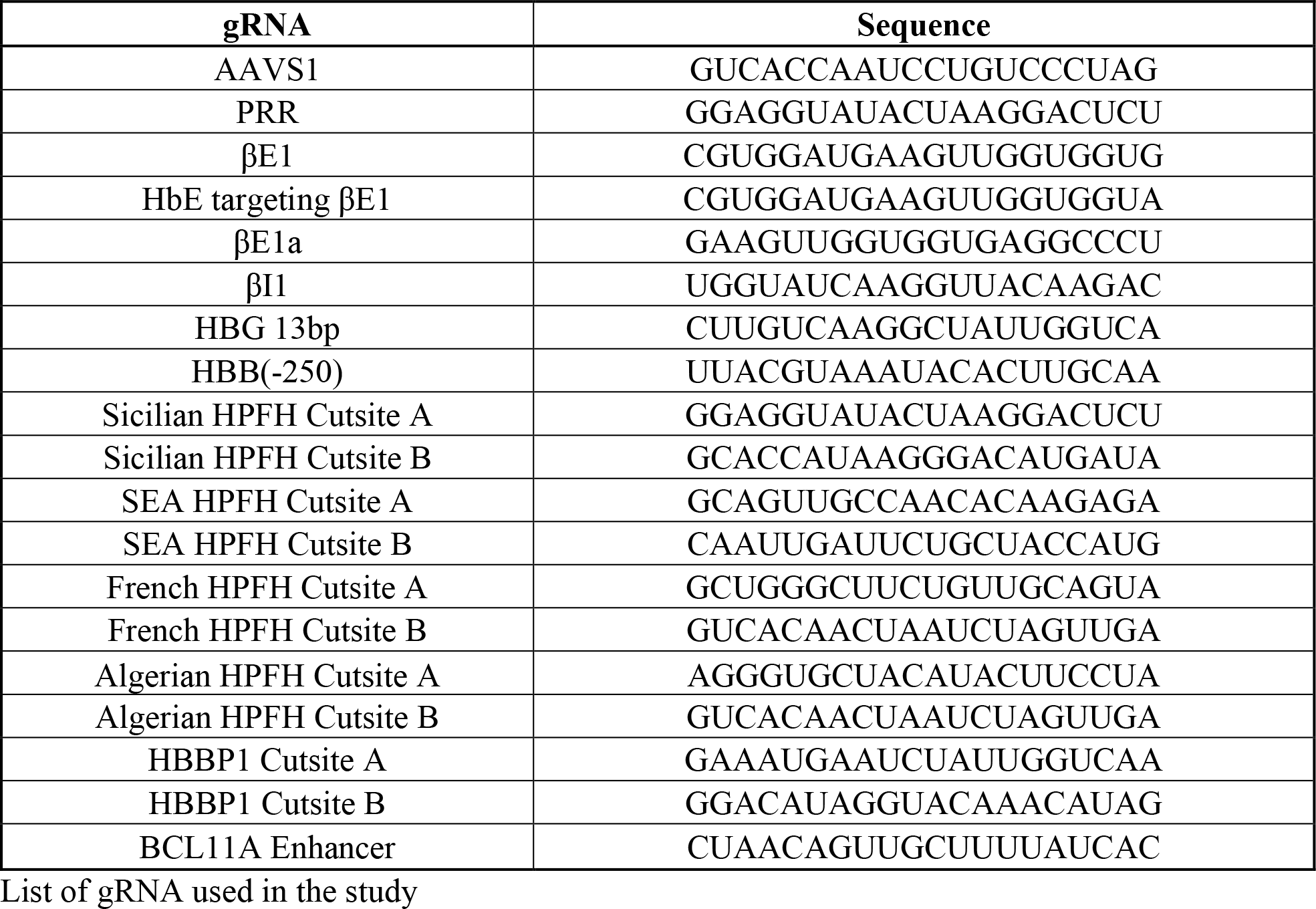

**Table – S3.**
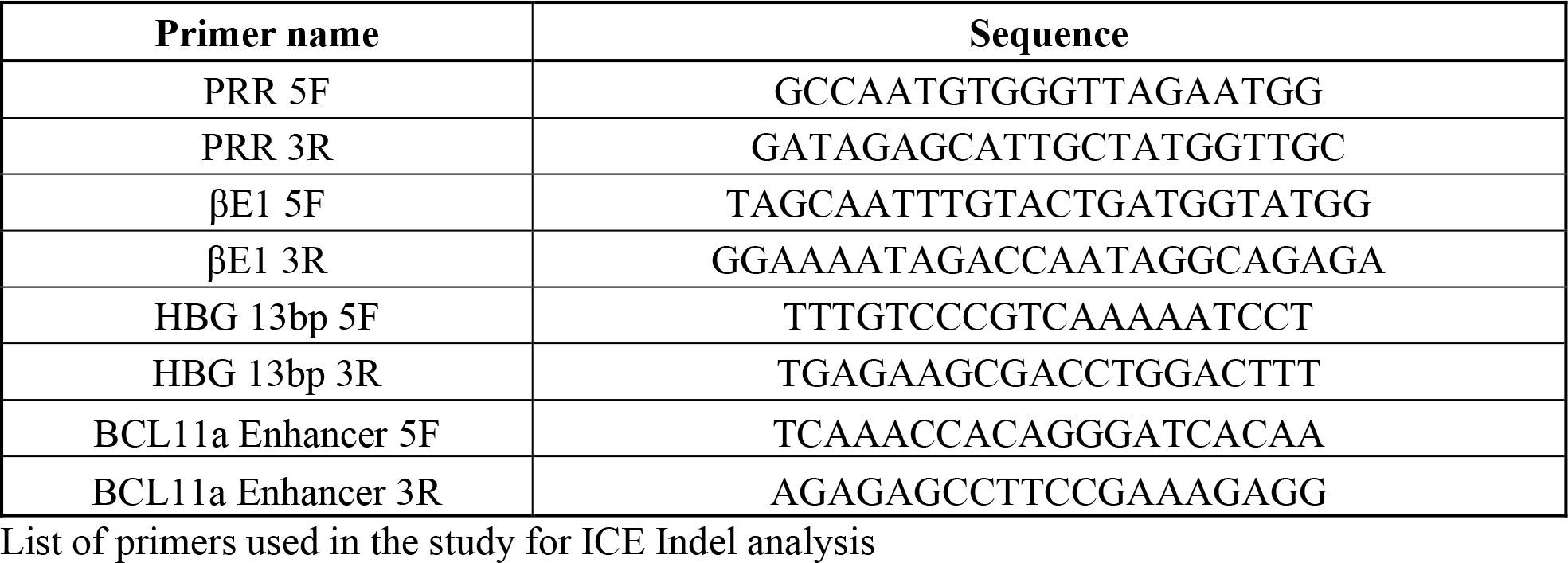

**Table – S4.**
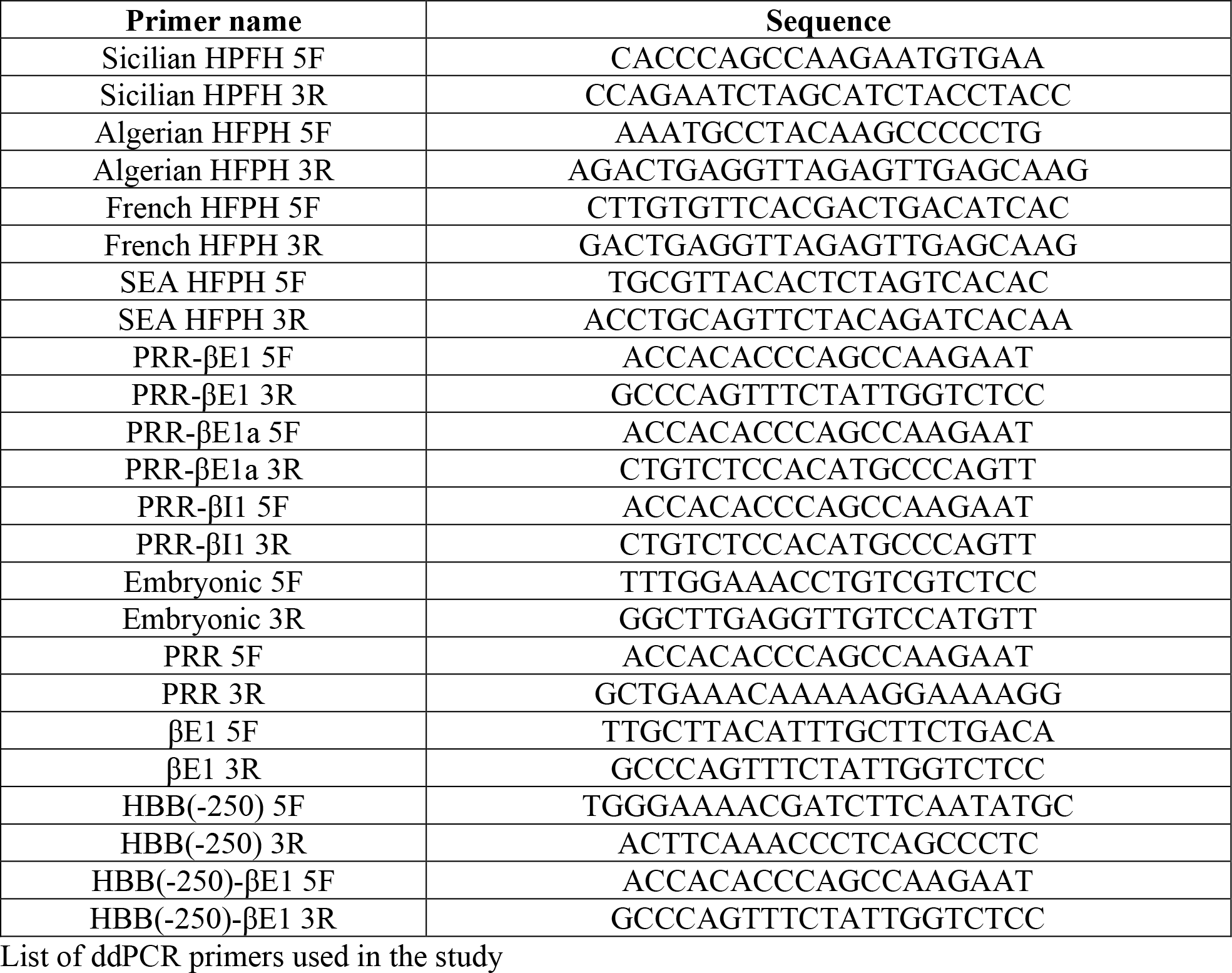

**Table – S5.**
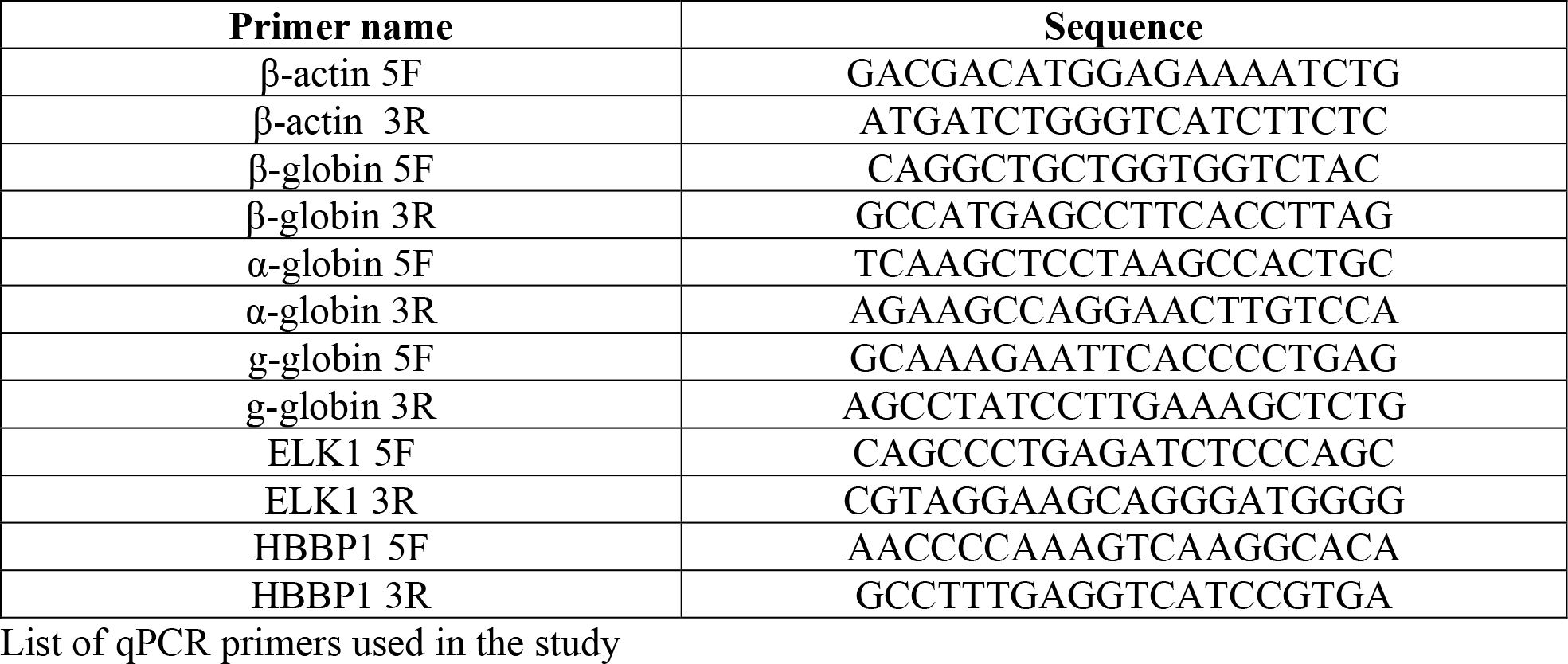

**Table – S6.**
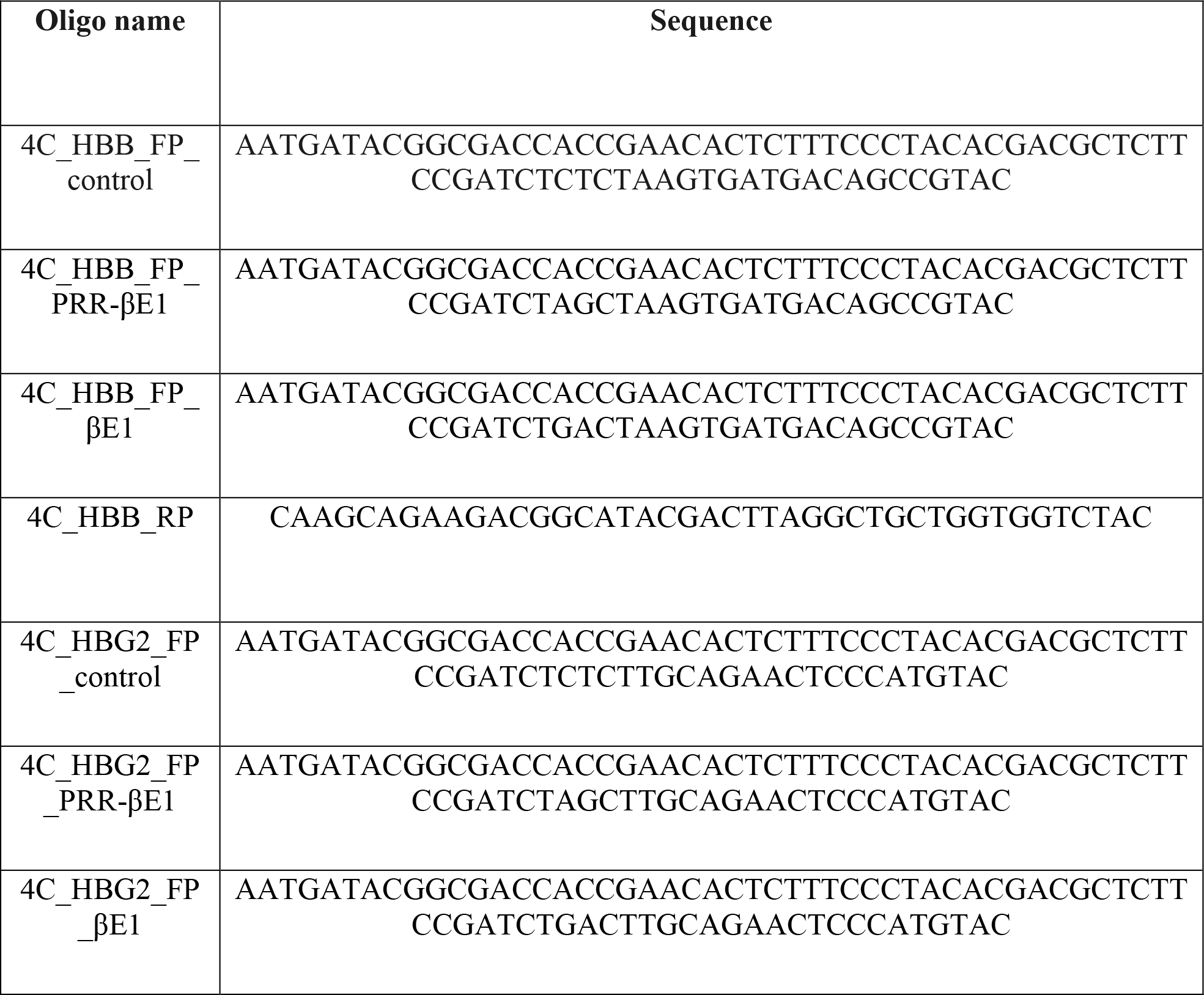
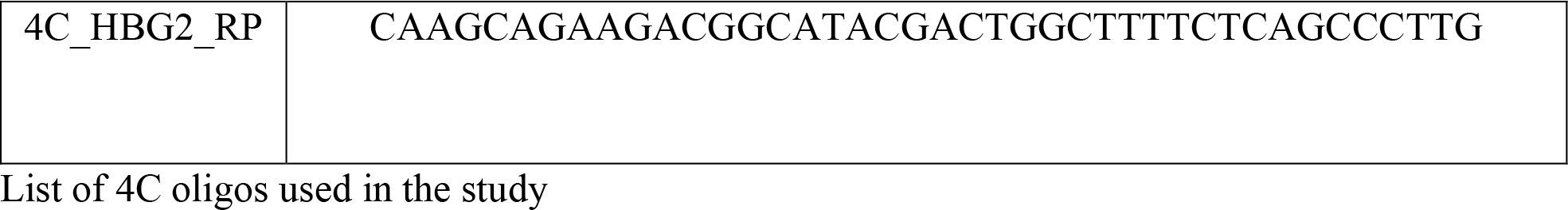

**Table – S7.**

| ANTIBODY/DYE | COMPANY | CAT NO |
| --- | --- | --- |
| Anti-Hemoglobin $\beta$ | Santa Cruz Biotechnology | sc-21757 |
| Anti-Hemoglobin $\alpha$ | Santa Cruz Biotechnology | sc-514378 |
| Anti-Hemoglobin $\gamma$ | Santa Cruz Biotechnology | sc-21756 |
| Anti-Actin Ab-5 | BD | 612656 |
| Goat anti-Mouse IgG | Invitrogen | 31430 |

List of western blot antibodies used in the study

